# Structural conservation and expanded functionality of hyper-stable human serum albumin variants

**DOI:** 10.64898/2026.04.10.717531

**Authors:** Sofia De Felice, Christian Buratto, Alessia Savio, Maria Morbidelli, Emanuele Papini, Laura Acquasaliente, Kristin Hovden Aaen, Jeannette Nilsen, Jan Terje Andersen, Alessandro Angelini, Arjen J Jakobi, Laura Cendron

## Abstract

Human serum albumin (hSA) is the most abundant protein in human plasma, and its pharmacological properties, such as long plasma half-life mediated by the neonatal Fc receptor (FcRn) and its ability to bind endogenous and exogenous molecules, make it attractive for biotechnological applications. Currently, most wild type (WT) SAs are derived from human or bovine serum or produced in yeast and mammalian cells. Although well established, these methods are costly, difficult to reproduce, and not environmentally sustainable. Building on a previous study to design highly mutated hSA sequences, we extend the validation through an in-depth analysis of three engineered hSA variants; hSA1, hSA2, and hSA3, containing 16, 25, or 73 amino acid substitutions, respectively. These variants were designed for enhanced solubility, stability, and expression in Escherichia coli. All three variants showed low- micromolar affinities for hFcRn at pH 5.5, and negligible binding at pH 7.4. In a human endothelial cell-based recycling assay (HERA), the engineered hSA variants were recycled by hFcRn to the same extent as hSA isolated from serum. Exploring the properties of canonical drug-binding sites, warfarin affinity was comparable to WT hSA, whereas ibuprofen binding differed. Complementary cytotoxicity assays on human macrophages confirmed negligible toxicity and biocompatibility. A cryo-electron microscopy structure of hSA3 revealed that, despite extensive engineering, the native heart-shape of hSA, folding of domains, and its open conformation were preserved. These findings validate the structural integrity and functional adaptability of engineered hSA variants, underscoring their potential as versatile, animal-free solutions for next-generation therapeutics and biotechnological applications.

## 1. Introduction

Human serum albumin (hSA) is the most abundant plasma protein in the human circulatory system, with concentrations ranging from 0.5 to 1 mM. The high concentration is maintained by constitutive production and a cellular recycling mechanism mediated by a pH-dependent interaction with the neonatal Fc receptor (FcRn) providing it with a plasma half-life of approximately 21 days [1–4]. Briefly, circulating hSA, which is taken up by endothelial cells lining a blood vessel by fluid-phase pinocytosis, may engage FcRn in acidified endosomes (pH < 6). Following binding, the complexes are sorted into vesicles that traffic back to the plasma membrane, where they undergo exocytosis. Exposure to the physiological pH of the blood (pH 7.4) triggers dissociation and release of hSA back into the blood. Proteins that do not bind FcRn are rather directed towards lysosomal degradation, and hence, albumin is saved from degradation, explaining its long plasma half-life [5–10].

The multifunctional roles of albumin make it a critical component in maintaining oncotic pressure, transporting an array of endogenous and exogenous molecules, and facilitating pharmacological interventions (Fig. 1A; [1–3]). Clinically, hSA derived from human plasma has been extensively used in the treatment of conditions such as hypovolemia, hypoalbuminemia, liver disease, nephrotic syndrome, and sepsis [2]. Originally introduced in the 1940s to prevent osmotic shock following hemorrhage, the therapeutic applications of hSA have broadened to include its use in the management of burn injuries, stabilization of vaccines, and correction of hypoproteinemia [11]. Beyond its therapeutic utility, hSA, alongside bovine serum albumin (bSA), serves as a fundamental component in pharmaceutical and biochemical research [12]. It is widely employed as a stabilizer and dispersant in drug delivery systems, a molecular weight standard in protein purification, and as an additive in biochemical assays such as enzyme-linked immunosorbent assays (ELISA) and Western blots (WB) [12]. In cell culture, albumin is pivotal in serum formulations, enhancing cell viability and proliferation [12]. More recently, recombinant hSA has gained attraction in the cultured meat industry, offering a viable alternative to fetal bovine serum (FBS) in the growth of bovine satellite cells [13]. As the global demand for recombinant proteins rises steeply across therapeutic, biotechnological, and industrial spheres, the dependence on animal-derived albumin remains widespread. This poses several challenges, including batch-to-batch variability, animal welfare concerns, and the risk of contamination with animal-derived pathogens and DNA [14]. In light of these concerns, recent research endeavors have shifted towards the development of engineered hSA variants designed to enhance structural stability, improve pharmacological efficacy, and offer application-specific functionality [15]. Generating such novel albumin derivatives hold significant potential to expand reproducibility, safety, and efficacy in many biomedical and industrial applications. Three engineered hSA variants were initially conceptualized *in silico*, employing the Protein Repair One Stop Shop (PROSS) algorithm. This algorithm leverages structural and sequence similarity to albumin from other species introducing strategic mutations aimed at improving solubility, stability, and expression levels in *Escherichia coli* [15]. Notably, the PROSS algorithm integrates atomistic design calculations with phylogenetic analysis, enabling the identification and incorporation of stabilizing mutations without compromising the native architecture and topology of the protein [16,17]. Prior to the advent of PROSS, the recombinant production of hSA was predominantly restricted to yeast, rice, mammalian cell lines, or directly purified from serum often resulting in prohibitively high costs of several hundred euros per milligram of protein.

**Fig. 1.**
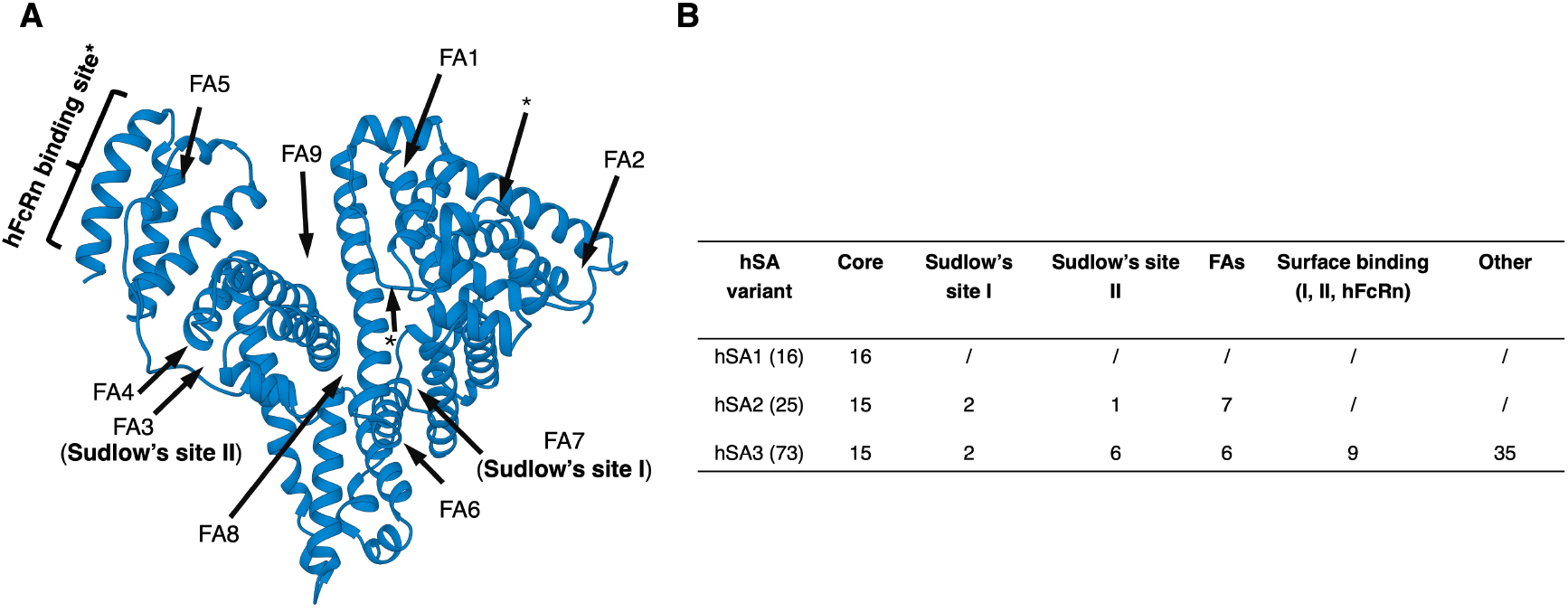
hSA variants analysis. (**A**) Three dimensional structure of hSA in the open conformation (PDB code 2BXI), fatty acids (FA), Sudlow’s I, II, and hFcRn binding sites are annotated. Classification and annotation of hSA1, hSA2 and hSA3, with the number of amino acid substitutions in each region indicated. (**B**) Amino acid substitutions located in regions that bind both fatty acids and drugs are classified as occurring in the drug binding sites.

In this study, we present a detailed comparative analysis on structure and function of three engineered hSA variants; hSA1 (16 mutations, the most conservative design with no surface mutations and no mutations in the binding sites), hSA2 (25 mutations, additional mutations in the binding sites), and hSA3 (73 mutations throughout the entire amino acid sequence; Fig. 1B). Among these engineered variants, hSA1 has undergone a preliminary characterization both structurally and functionally [15]. The structure of hSA1 was determined through X-ray crystallography at a resolution of 2 Å, revealing no significant differences compared to WT hSA in its open conformation (PDB ID 8A9Q; [15]). Notably, electron density for lipid molecules was detected in the crystal structure of hSA1, even in the absence of added lipids during crystallization, suggesting an interaction with myristic acid derived from the bacterial cultures [15]. Additionally, the stabilizing mutations incorporated within the core of hSA1 resulted in enhanced binding affinities for the ligands warfarin and ketoprofen, exceeding those of commercial WT hSA [15].

In the present study, we have studied the human FcRn (hFcRn) binding properties of hSA1, hSA2, and hSA3, by surface plasmon resonance (SPR). The binding kinetic measurements were complemented by a human endothelial cell-based recycling assay (HERA), which is a cellular assay used to evaluate FcRn-dependent recycling of albumin-based therapeutics [18–21]. Furthermore, the impact of mutations on binding affinities towards canonical hSA ligands, such as ibuprofen and warfarin, was investigated. Cytotoxicity assays performed on primary human monocyte–derived macrophages were employed to evaluate the biocompatibility of the designs. Finally, a cryo-electron microscopy (cryo-EM) structure of hSA3 was obtained to evaluate the three-dimensional folding of the most extensively mutated variant. These results provide insight into the design space available for developing functional, non-animal-derived albumin variants for biotechnological applications.

## 2. Results and Discussion

### 2.1 Production, Thermal stability and Dynamic light scattering measurements

The three engineered hSA variants, named hSA1, hSA2, and hSA3, were isolated in milligram quantities from *E. coli* cultures following the procedure adapted from Kersonsky *et al*. [8]. An additional thermal treatment was introduced in the purification steps for all the variants, significantly improving the purity of the isolated samples (Fig. S1, S2, S3A). Yields, solubility and thermostability followed the expected trend of properties improving with the number of stabilizing mutations, and reaching, in the case of hSA3, hundreds of milligrams per liter of culture and a Tm above 100 °C, as reported in literature [15].

To assess the dispersity and particle size of the purified hSA variants, dynamic light scattering measurements were conducted on the purified proteins at a concentration of 5 mg/mL (Fig. 2A). All variants displayed mean sizes and polydispersity indexes (PDI) comparable to those of commercial defatted hSA, with superimposable aggregation profiles (Fig. 2B and Fig. S3B-F).

**Fig. 2.**
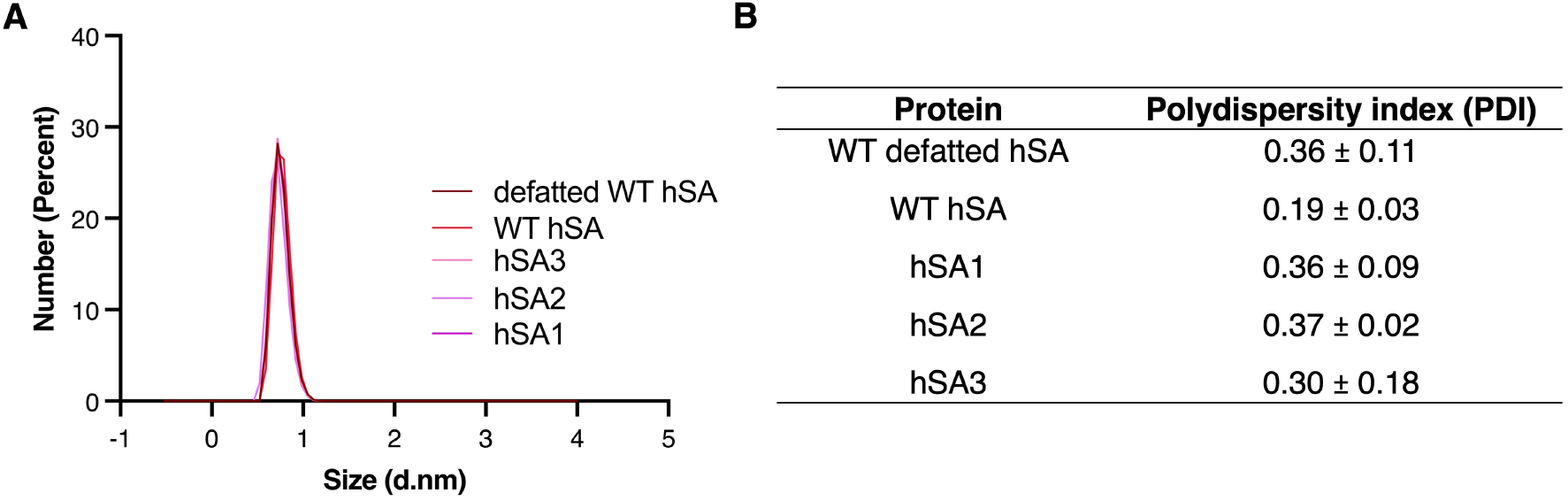
Dynamic light scattering analysis of defatted WT hSA, WT hSA, hSA1, hSA2, and hSA3. A Malvern Panalytical instrument was used to measure dynamic light scattering of hSA variants at 5 mg/mL in PBS (pH 7.4). (**A**) Hydrodynamic size distributions and (**B**) corresponding polydispersity index (PDI) values are shown as the mean ± standard deviation of three measurements with five replicates in each.

### 2.2 Binding of gold-standard albumin drug ligands to engineered hSA variants

hSA binds several endogenous and exogenous molecules, including warfarin and ibuprofen. To evaluate the binding capacities of hSA2 and hSA3 to these gold-standard hSA ligands, isothermal titration calorimetry was performed (ITC; Fig. 3, 4, S4, S5).

**Fig. 3.**
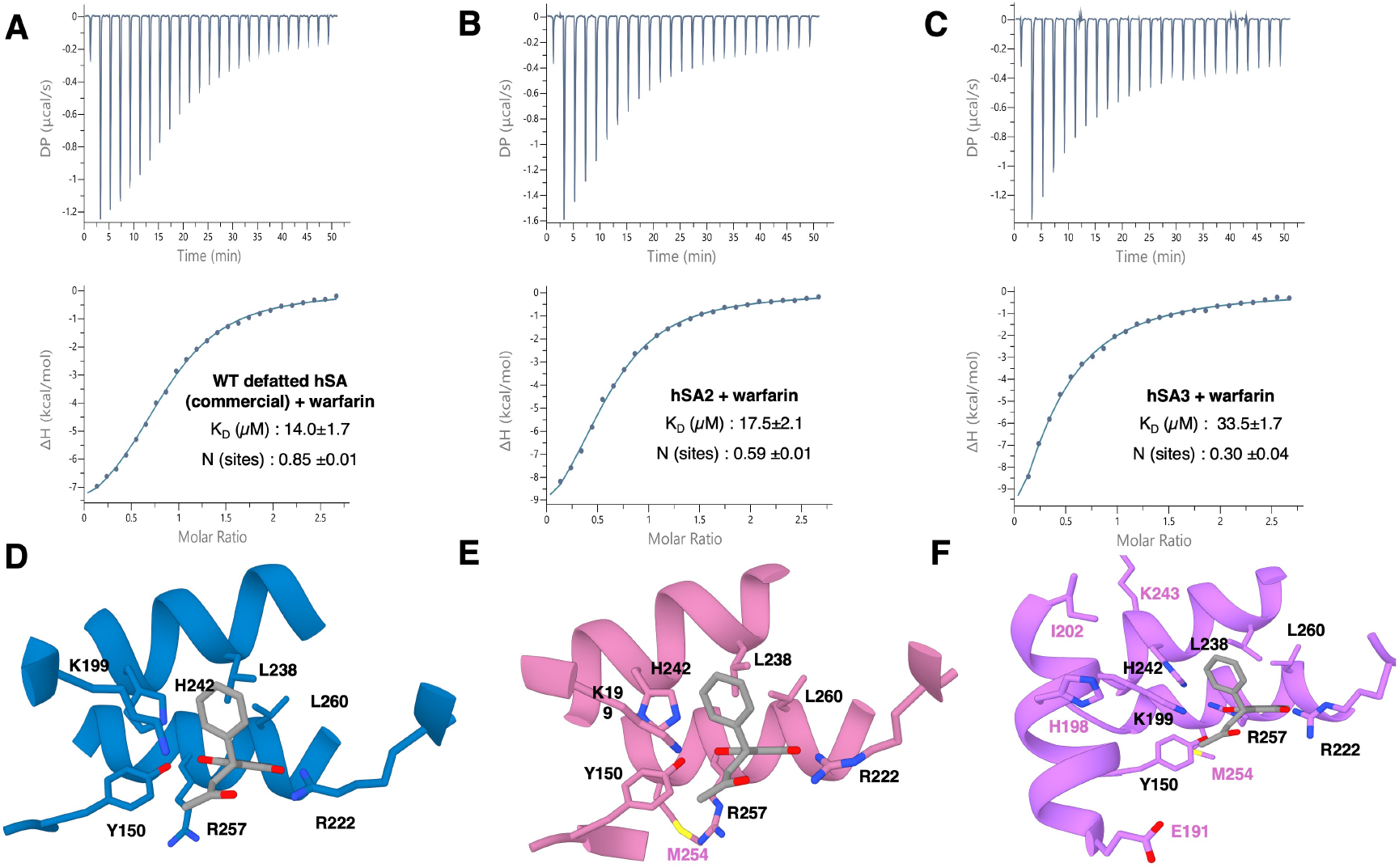
Warfarin binding to hSA variants measured by ITC. Binding is shown for (**A**) commercial defatted WT hSA, (**B**) hSA2, and (**C**) hSA3. (**D**) Sudlow site I of WT hSA in complex with warfarin (PDB code 2BXD) is shown, with key binding residues annotated. Amino acid substitutions near Sudlow site I of (**E**) hSA2 (modeled with Alphafold3) and (**F**) hSA3 (PDB code 9R8P) are annotated in magenta. In both hSA2 and hSA3, warfarin was docked in the same position as observed in WT hSA. The reported K_D_ values and number of binding sites represent the mean of three independent replicates.

**Fig. 4.**
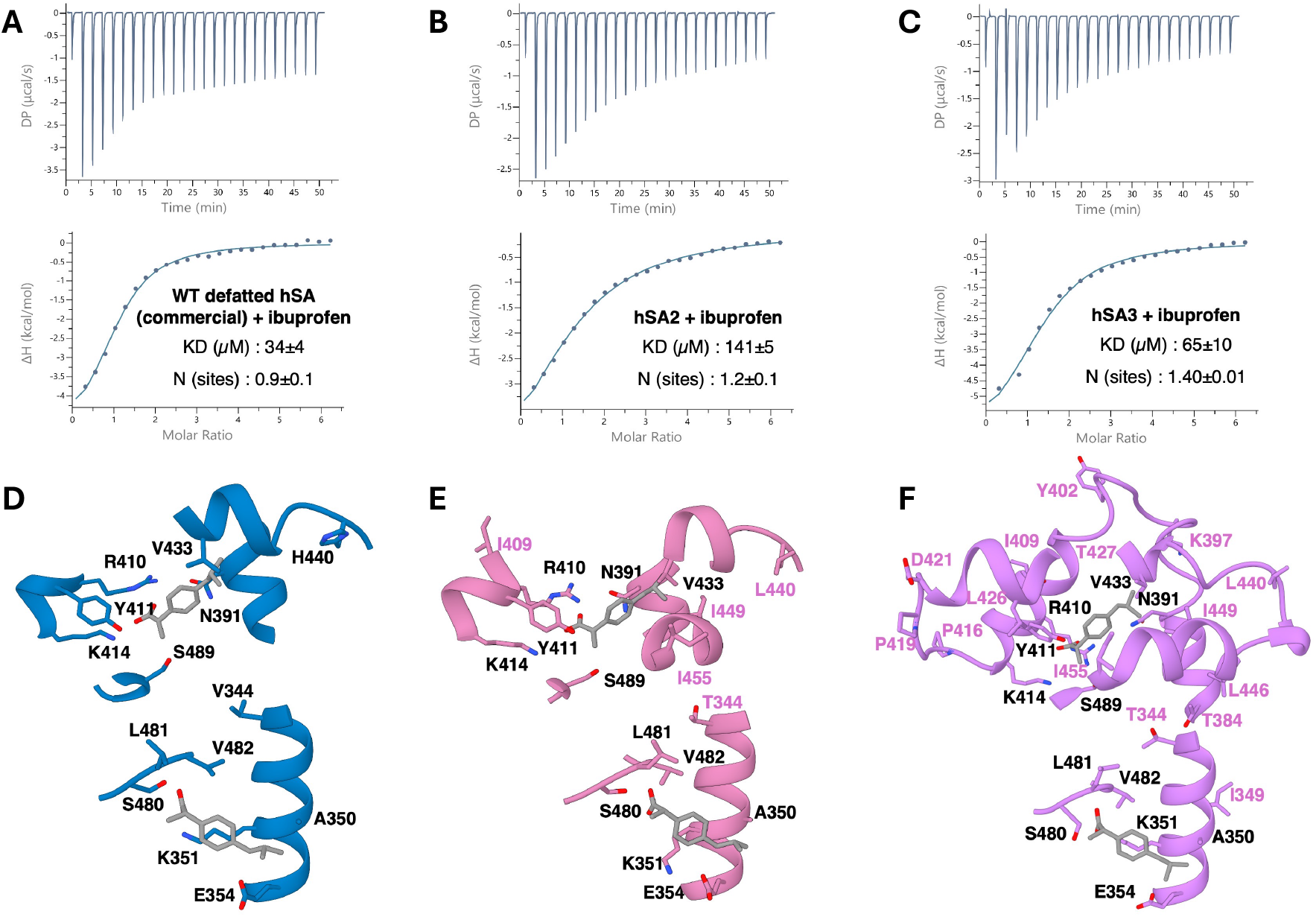
Ibuprofen binding to hSA variants measured by ITC. Binding is shown for (**A**) commercial defatted WT hSA, (**B**) hSA2, and (**C**) hSA3. (**D**) The crystal structure of WT hSA in complex with ibuprofen at Sudlow site II (PDB code 2BXG) is shown, with key binding residues annotated. Mutations near the binding sites in (**E**) hSA2 (modeled with Alphafold3) and (**F**) hSA3 (PDB code 9R8P) are annotated. In both hSA2 and hSA3, ibuprofen was docked in the same position as done for WT hSA. The reported K_D_ values and number of binding sites represent the mean of three independent replicates.

The ITC experimental binding curves demonstrated that hSA2 and hSA3 retained binding affinity for warfarin. Despite the introduction of mutations in the warfarin-binding pockets, hSA2 bound warfarin with an affinity in the same order of magnitude as that obtained for both commercial WT hSA and defatted WT hSA (Fig. 3A-B, S4; Table 1). hSA3 showed decreased affinity for warfarin, with a dissociation constant (K_D_) roughly twice that of hSA2 (Fig. 3B; Table 1; Table S1). Importantly, the key amino acid residues that form the hydrophobic pocket where warfarin binds in its main binding site remain unchanged in hSA2 and hSA3, and thus, the observed change in affinity for hSA3 cannot be attributed to the mutations introduced in Sudlow site I of this variant. Analogously, the key polar residues such as R222, R257 or H242, well-known for their capacity to interact with the phenoxide group of warfarin by ion-pairing are conserved (Fig. 3D-F; [22]). The few notable mutations adjacent to the warfarin-binding hydrophobic pocket are A191E and T243K. Although they do not play a direct role in interacting with warfarin, we speculate that T243K could disfavor the function of the adjacent H242 by shifting its acid-base equilibrium. Moreover, the 73 mutations present in the hSA3 variant could have introduced epistatic and long-distance effects that, although indirect, could influence the affinity properties of hSA3 for any ligand. Such a large number of mutations could have altered the plasticity of the protein and thus the necessary adaptation of the cavity-defining helices, such as α-IV of domain IB (from Y150 to C169), α-IV of domain II (from P281 to A291) or α-III of domain IIA (from D249 to K262), ultimately affecting the features of the cavity and/or the positioning of key residues in the binding pocket (Y150, R257; Fig. 3D,3F).

**Table 1.** Ligands binding to hSA variants measured by ITC. The values reported are the average of three replicates and the standard deviation (s.d) is reported. The symbol “—” is used to report a not interpretable binding profile.

| Drug | hSA variant | K<sub>D</sub> mean (μM) | Number of sites |
| --- | --- | --- | --- |
| Warfarin | Defatted WT hSA | 14.0 ± 1.7 | 0.85 ± 0.01 |
|  | WT hSA | 13.4 ± 1.7 | 0.60 ± 0.02 |
|  | hSA2 | 17.5 ± 2.1 | 0.59 ± 0.01 |
|  | hSA3 | 33.5 ± 1.7 | 0.30 ± 0.04 |
| Ibuprofen | Defatted WT hSA | 33.9 ± 3.7 | 0.9 ± 0.1 |
|  | WT hSA | — | — |
|  | hSA2 | 141 ± 5 | 1.2 ± 0.1 |
|  | hSA3 | 65 ± 10 | 1.40 ± 0.01 |

To further investigate the impact of the introduced mutations on the binding capacity of Sudlow site II, one of the principal drug-binding *loci* alongside Sudlow site I, we measured binding to ibuprofen. In contrast to the affinity reported for hSA1 by Khersonsky and colleagues, hSA2 exhibited a four-fold reduction in ibuprofen affinity relative to commercial defatted WT hSA (Fig. 4A-B; Table 1; [8]). This may be ascribed to the mutations introduced in the region close to the ibuprofen binding pocket (Fig. 4E; Table S2). For example, the mutation V344T may have altered some hydrophobic interactions, and the bulky A449I may have caused ibuprofen repositioning to avoid clashes with its side chain (Fig. 4E; Table 1). Besides, the mutation H440L may have reduced the electrostatic contribution involved in ibuprofen engagement (Fig. 4E; Table 1). Here, the comparison was done with commercial defatted WT hSA, since the standard hSA behavior has been proven to be affected by a partial loading of fatty acids. Indeed, fatty acids binding is known to interfere with ibuprofen binding affinity, since Sudlow site II is a preferential site for lipids that occupy the well-characterized FA4 and FA3 positions (Fig. 1; [1,23,24]).

Surprisingly, hSA3 showed slightly recovered binding to ibuprofen, having only two-fold weaker than that of defatted WT hSA (Fig. 4C). Such behaviour could be explained by the impact of epistatic effects introduced by the 73 mutations. Indeed, several residues in the region near Sudlow site II were mutated to non-polar amino acids. These changes may be responsible for additional hydrophobicity, which might compensate for the removal of other interactions present in the WT protein, or a different capacity to load lipids (Fig. 4F). Overall, the K_D_ values measured are in the mid-micromolar range (Table 1), which would suggest negligible binding at all for proteins present in the serum at nanomolar concentrations. On the other hand, these values acquire relevance when considering the high physiological concentration of hSA [1]. Accurate interpretation of ligand K_D_ for hSA necessitates not only accounting for its physiological serum concentrations (0.5–1 mM) but also its intrinsic promiscuity in lipid and endogenous ligands binding, frequently involving shared binding pockets. The designed hSA variants characterized herein exhibit enhanced conformational stability, as demonstrated by Khersonsky and colleagues, and modulated ligand-binding profiles, enabling the tuning of their pharmacokinetic parameters [15].

### 2.3 Mutational remodeling of hSA variants does not affect hFcRn interaction

To assess whether the extensive mutational remodeling of hSA1, hSA2, and hSA3 affected the interaction with hFcRn, we performed surface plasmon resonance (SPR), where the hSA variants were injected over immobilized receptor. To mimic the pH-gradient of the trafficking endosomes during hFcRn-mediated cellular recycling of hSA, the process ensuring its long plasma half-life, the experiments were performed at pH 5.5, 6.0, and 7.4 (Fig. 5).

**Fig. 5.**
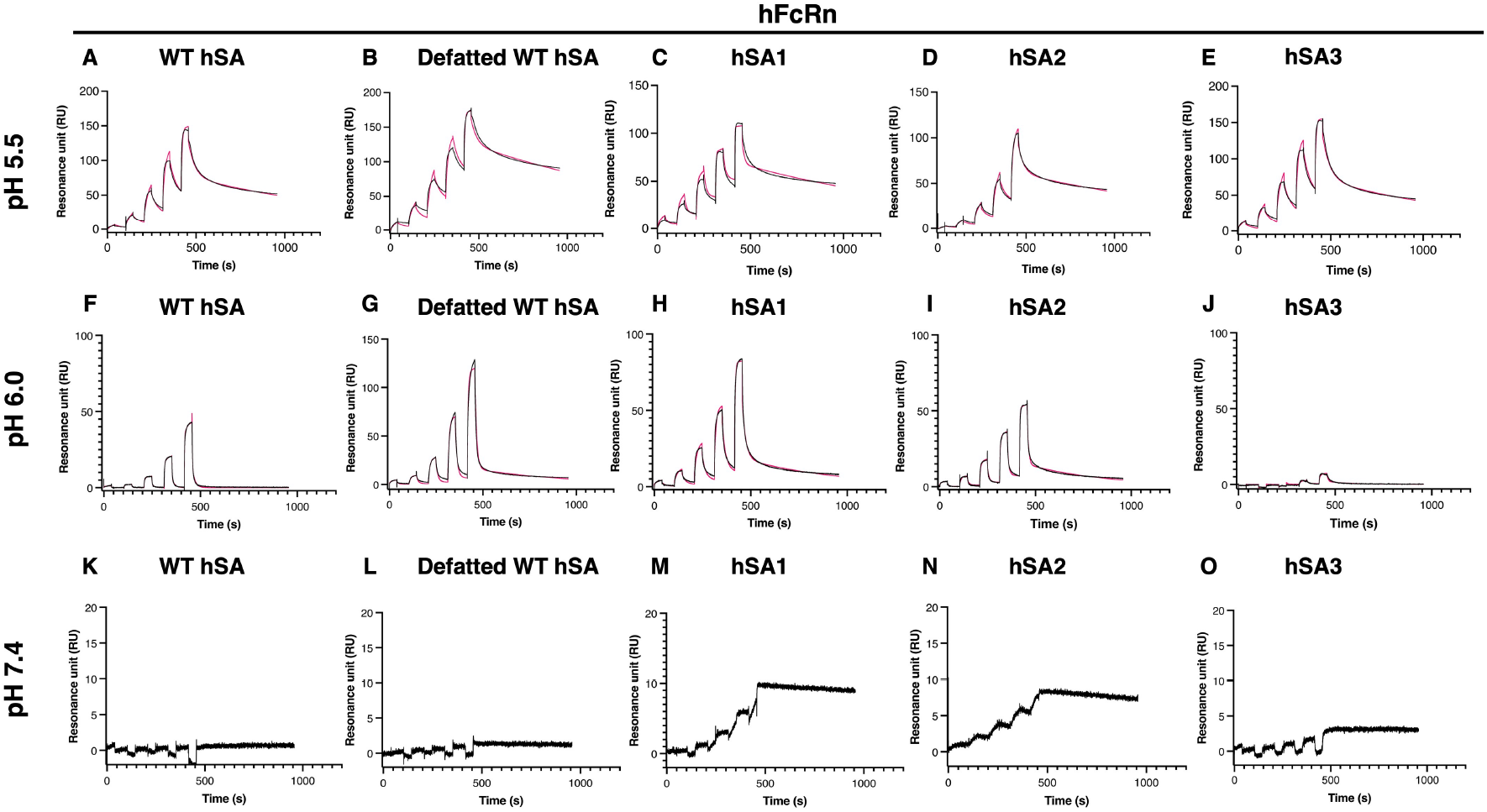
Representative single-cycle binding kinetics curves fitted using a two-state reaction binding model at pH 5.5, 6.0 and 7.4. Titrated concentrations (0.37, 1.1,3.3,10, 30 µM) of commercial WT hSA (**A, F, K**), commercial defatted WT hSA (**B, G, L**), hSA1(**C, H, M**), hSA2 (**D, I, N**), and hSA3 (**E, J, O**) were injected over immobilized recombinant hFcRn. Fitted curves are shown in red. Curves at pH 7.4 (**K-O**) were not fitted due to their low RU values.

The SPR experiments at pH 5.5, showed that the receptor binding affinity was preserved for all variants, indicating that the introduced mutations do not disrupt this interaction (Table 2, Fig. 5A-E, S6). The K_D_ values measured for the interaction between WT hSA and hFcRn were in line with previous reports [14]. Interestingly, hSA1 displayed a slightly higher affinity for hFcRn, showing a K_D_ of 0.22 µM compared to 0.8 µM for WT hSA, 1.1 µM for hSA2, and 0.9 µM for hSA3, due to an increased association rate constant (k_a_; Table 2, Fig. 5).

**Table 2.** SPR-derived kinetic values for binding of hSA variants towards hFcRn. The values reported are the average of three replicates and the standard deviation (s.d) is reported (Fig. S6, S7, Table S3). Albumin concentrations ranged from 0.3 to 30 µM and sensorgrams were globally fitted using the two-state reaction model (Biacore T100 Evaluation Software).

| pH | hSA variant | $K_{a1}$ ($10^4$ M<sup>-1</sup> s<sup>-1</sup>) | $K_{d1}$ ($10^{-3}$ s<sup>-1</sup>) | $K_{a2}$ ($10^{-3}$ s<sup>-1</sup>) | $K_{d2}$ ($10^{-3}$ s<sup>-1</sup>) | $K_D$ (μM) |
| --- | --- | --- | --- | --- | --- | --- |
| 5.5 | Defatted WT hSA | 2.1 ± 1.6 | 86 ± 70 | 15 ± 15 | 1.0 ± 0.1 | 0.3 ± 0.1 |
| | WT hSA | 0.6 ± 0.6 | 49 ± 50 | 12 ± 4 | 1.2 ± 0.3 | 0.8 ± 0.5 |
| | hSA1 | 2.8 ± 0.5 | 87 ± 14 | 12.0 ± 1.5 | 0.9 ± 0.1 | 0.22 ± 0.05 |
| | hSA2 | 0.20 ± 0.03 | 23.0 ± 3.3 | 9.7 ± 0.8 | 1.1 ± 0.1 | 1.10 ± 0.03 |
| | hSA3 | 0.6 ± 0.1 | 21.7 ± 2.1 | 5.3 ± 0.9 | 1.5 ± 0.6 | 0.9 ± 0.4 |
| 6.0 | Defatted WT hSA | 1.1 ± 1.2 | 295 ± 33 | 2.0 ± 0.1 | 2.2 ± 0.1 | 14.0 ± 1.5 |
| | WT hSA | 0.7 ± 0.1 | 540 ± 34 | 23 ± 4 | 76 ± 7 | 53 ± 1 |
| | hSA1 | 0.7 ± 0.1 | 58 ± 5 | 4.10 ± 0.01 | 2.4 ± 0.2 | 3.1 ± 0.7 |
| | hSA2 | 1.10 ± 0.02 | 144 ± 4 | 4.8 ± 0.1 | 2.6 ± 0.1 | 4.7 ± 0.4 |
| | hSA3 | 0.3 ± 0.1 | 136 ± 17 | 5.6 ± 1.1 | 5.5 ± 2.2 | 20 ± 9 |

When performing the same SPR setup at pH 6.0, all hSA variants showed reduced binding affinity to hFcRn compared to that measured at pH 5.5 (Table 2, Fig. 5F-J, S7). The binding of hSA1 and hSA2 was less affected by increasing the pH to 6.0 than WT hSA, indicating that, within the physiological context of the endosomal environment, these variants may be capable of engaging hFcRn at an earlier stage than WT hSA. This interpretation is consistent with the behaviour previously reported for mouse SA (Nilsen et al., 2018). To evaluate further how the mutations impact the affinity of hSA1, hSA2, and hSA3 towards hFcRn at physiological pH, where binding is not expected to occur, SPR was finally performed at pH 7.4. The sensorgrams showed no binding for all variants, including WT hSA and defatted WT hSA (Fig. 5K-O, S8), confirming that the expected pH-dependency of the interaction between hFcRn and hSA was preserved despite engineering.

Multiple sequence alignment (ClustalW) revealed that the hSA variants harbour mutations in hSA domains I and III, regions reported to mediate hFcRn binding (Fig. 6, [15,18]). Among the key residues present in domain III, three histidine residues (^hSA^H464, ^hSA^H510, and ^hSA^H535) are known to play a role in pH sensing, where protonation facilitates the interaction with hFcRn [Fig. 6A, [15]. Nuclear magnetic resonance (NMR) studies have shown that at pH lower than 6.0, ^hSA^H510 and ^hSA^H535 in domain III are protonated, introducing positive charges and facilitating hFcRn binding [15]. Previous studies have demonstrated that the replacement of ^hSA^H464, ^hSA^H510 and ^hSA^H535 with a glutamine or a phenylalanine abolishes or greatly reduces hFcRn binding affinity, underscoring their functional importance [25,26]. ^hSA^H510, which establishes pH-dependent interactions with ^hFcRn^W176, along with ^hSA^H435 and ^hSA^H535, are conserved across all hSA variants. On the contrary, ^hSA^H440 is replaced with a leucine in hSA2 and hSA3 with no major impact on hFcRn engagement as confirmed by SPR and HERA (see below). This is in line with a previous study showing that hFcRn binding was unaffected by substitution of ^hSA^H440 with a glutamine [26]. The alignment of hSA variants further confirmed the conservation of residues within the H-H loop that interact with ^hFcRn^W53 (*i*.*e*., F507, T508, F509), ^hFcRn^W59 (*i*.*e*., T467, L463, H464, L460) and β_2_ -microglobulin, as well as the hSA C-terminal alpha helix, in particular the residue ^hSA^L585, whose truncation has been shown to decrease hFcRn affinity ([19,27]; Fig. S6).

**Fig. 6.**
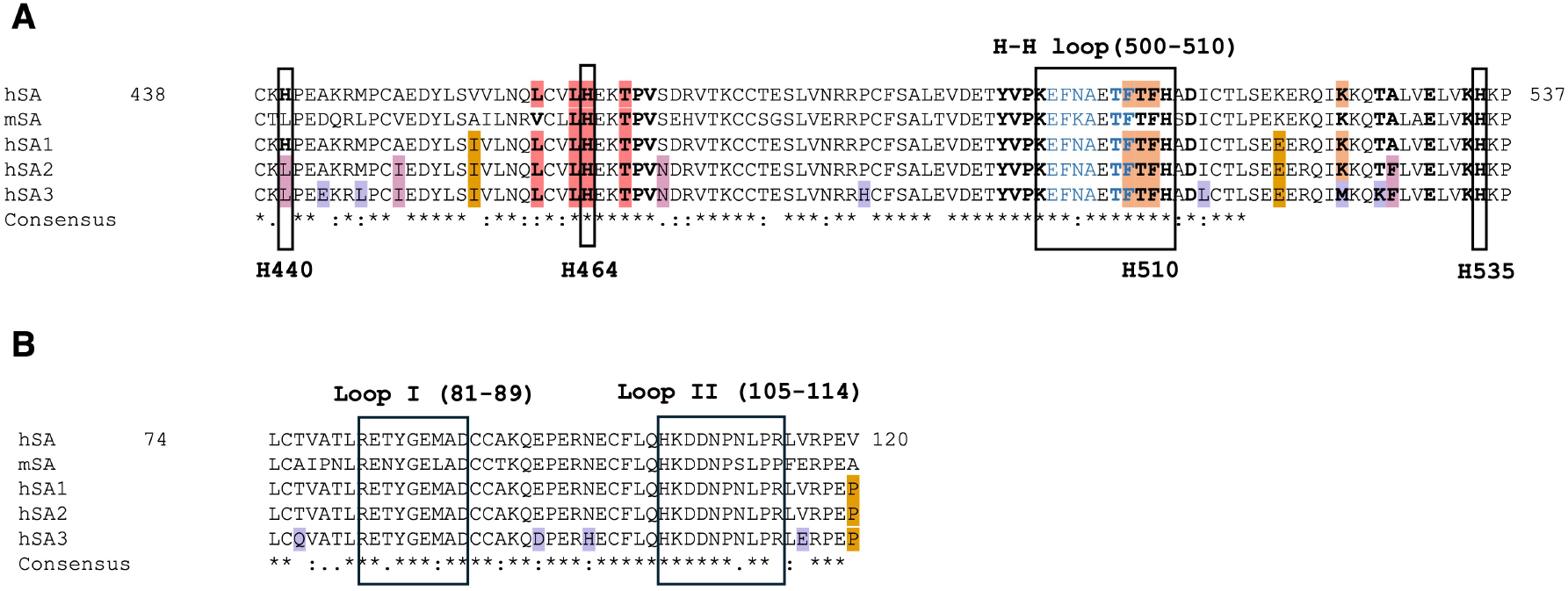
Multiple sequence alignment of WT hSA, mSA, hSA1, hSA2 and hSA3 regions involved in hFcRn binding. Multiple amino acid sequence alignments of hSA from serum, mSA, hSA1, hSA2, and hSA3performed using the ClustalW. (**A**,**B**) Alignment of amino acid residues 438-537. (**A**) Amino acid substitutions introduced in hSA1 (gold), hSA2 (pink), and hSA3 (violet) are indicated. Histidines essential for pH-dependent hFcRn binding (H440, H464, H510, H535) are boxed. Residues involved in hFcRn binding are in bold, those interacting with β2-microglobulin are in dark blue, while residues interacting with both hFcRn and β2-microglobulin are in bold dark blue. Residues interacting with ^FcRn^W53 are highlighted in orange, and those interacting with ^FcRn^W59 in red. (**B**) Alignment of amino acid residues 74-120 of domain I-, using the same color code as in panel A is used. Residues involved in FcRn binding are boxed. Loop I and II, involved in hFcRn binding are boxed.

Nearby regions of hSA3 hold permissive mutations such as ^hSA^K524M and ^hSA^T527K, as well as ^hSA^A528F, the latter present in hSA2 too. Notably, ^hSA^K573 residue, which is mutated to serine in hSA3, was previously reported to enhance hFcRn affinity by 5- and 17-fold when replaced with serine and proline, respectively [27,28]. Further permissive substitutions are observed in the flanking regions of loop I (R81-D89) and loop II (^hSA^H105-R114) of domain I, including ^hSA^T76Q, ^hSA^E95D, ^hSA^N99H, ^hSA^V116E, and ^hSA^V120P (Fig. 6B).

### 2.4 HERA assay proves of hFcRn-mediated cellular recycling in human endothelial cells

To study hFcRn-mediated recycling of hSA1, hSA2, and hSA3 in an animal-free system, HERA was performed [21,29]. Here, human endothelial cells stably expressing hFcRn (HMEC-1 hFcRn) were seeded in 48-well plates. The following day, the cells were incubated for four hours with one of the three engineered hSA variants or commercial hSA isolated from serum. In addition, recombinant WT hSA and hSA QMP, an engineered variant that binds favourably to hFcRn, were included as controls in the assay, confirming that the assay was able to distinguish between hSA variants displaying different hFcRn binding affinities ([19,30]; Fig. S9). Next, the cells were lysed to enable quantification of the protein taken up by the cells. A parallel plate was treated identically, however, instead of lysing the cells after the four-hour incubation, serum-free medium was added. The next day, the cell medium was collected for quantification of the amount of protein recycled and released from the cells. Protein quantification in the cell lysate and medium was performed by ELISA.

All three engineered hSA variants were taken up by the endothelial cells to a similar extent as hSA isolated from serum (Fig. 7A, Fig. S10). Furthermore, while hSA1 and hSA2 were recycled equally well as that of albumin from human serum, about 0.73-fold less of hSA3 was detected in the recycling media (Fig. 7B, Fig. S10). Notably, these observations may be a result of differences in binding kinetics, as revealed by the SPR measurements at pH 6.0, which suggest that hSA1 and hSA2 may engage hFcRn at an earlier time point in the endosomal pathway than hSA3. Further studies in animal models are needed to determine how the amino acid substitutions affect the pharmacokinetic properties of albumin.

**Fig. 7.**
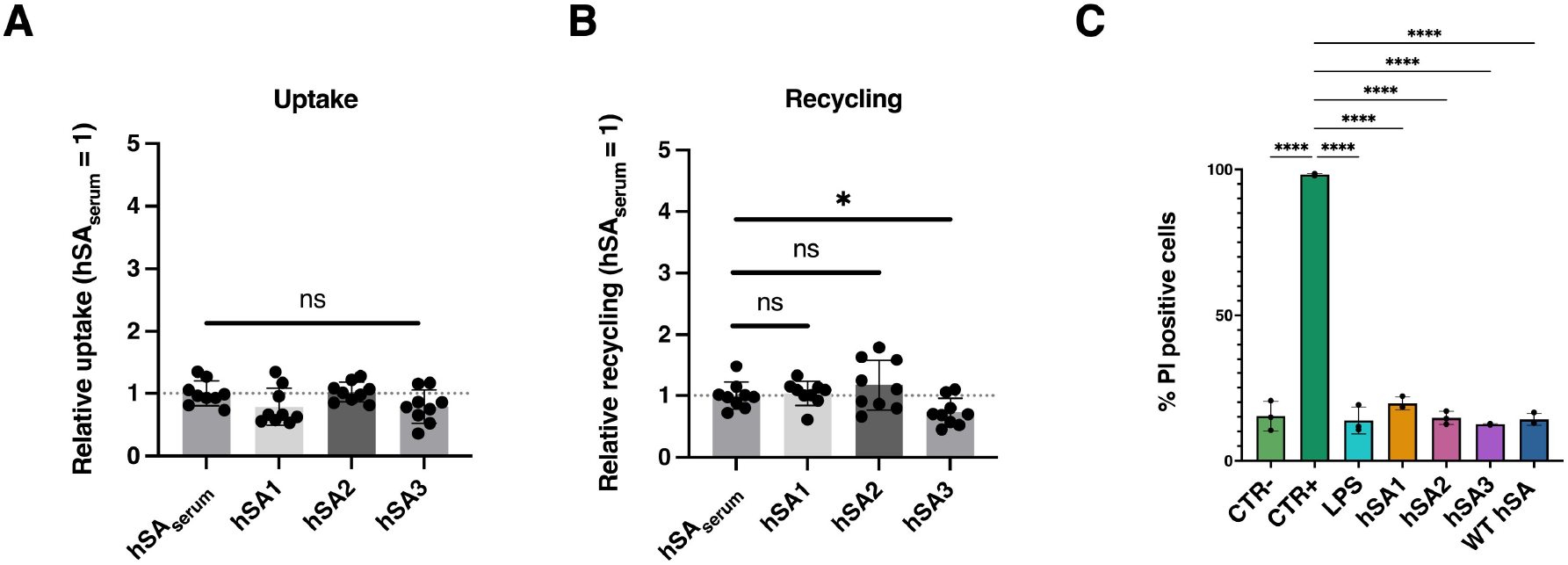
hFcRn-mediated cellular recycling and evaluation of biocompatibility of hSA variants. (**A, B**) Results from HERA with human endothelial cells stably overexpressing hFcRn showing cellular uptake of hSA variants (**A**) and hFcRn-mediated cellular recycling of hSA variants (**B**). Statistical significance was evaluated by Student’s t tests (* p = 0.0205). Data are shown as relative to that of hSA isolated from serum (hSA_serum_ = 1) and presented as the mean ± standard deviation (n = 9). (**C**) Results from biocompatibility assay with macrophages showing the percentage of PI-positive cells after exposure to hSA variants or WT hSA. CTR-, untreated macrophages; CTR+, cells treated with necrosis-inducing Ludox. Statistical significance was evaluated by one-way ANOVA (**** p < 0.0001).

### 2.5 hSA variants are biocompatible with human monocyte-derived macrophages

To assess the biocompatibility of the engineered hSA variants, we conducted propidium iodide (PI) staining on human monocyte-derived macrophages exposed to each variant at a concentration of 0.4 mg/mL, none of the hSA variants induced significant levels of necrosis (Fig. 7C). The quantification of PI-positive cells (background-subtracted) showed non-significant differences between cells treated with hSA variants and the untreated control, as determined by one-way ANOVA (Fig. 7C). These results suggest that, at the tested concentration, the hSA variants do not elicit cytotoxic effects on macrophages. Although preliminary, this assay supports the overall biocompatibility of the engineered hSA variants and lays the groundwork for future studies aimed at evaluating their potential immunogenicity.

### 2.6 The hSA3 variant adopts the open conformation of WT hSA

Due to the high solubility of the hSA3 variant, all attempts to obtain crystals for X-ray diffraction were unsuccessful. As an alternative, cryo-EM was employed to evaluate the structural consequences of the 73 introduced mutations on the overall fold of hSA3. To increase its size and randomize particle distribution in ice, hSA3 was complexed with a megabody construct (hereinafter referred to as MgbAlb1c7HopQ) previously characterized in complex with WT hSA by X-ray crystallography [31,32]. MgbAlb1c7HopQ comprises a nanobody (NbAlb1) that is cross-reactive to both mouse and human SA, fused to the circularly permutated extracellular domain of the adhesin HopQ of *Helicobacter pylori* [31].

The structure of hSA3 in complex with MgbAlb1c7HopQ was obtained by cryo-EM at an overall resolution of 3.9 Å (Fig. 8A-8B, S10-S11; Table S4; PDB ID 9R8P, EMD-53835), with local resolution ranging from 3.5 Å in the core to 8 Å at the periphery (Fig. S11). Residues from 3 to 583 were placeable in the density map, while the N-terminal His-tag was not visible, as expected. The structure of hSA3 presented the classical albumin heart-shape and the organization of the three homologous domains I, II and III (Fig. 8A), as already extensively described in literature for WT hSA [1–3]. Due to the low resolution of the extra density corresponding to NbAlb1, which protrudes from hSA3 domain II, NbAlb1 could not be modelled (Fig. 8B-8E-F). Additionally, the c7HopQ scaffold domain was not visible, likely due to its flexibility, a feature already observed in the cryo-EM structure of other complexes using the megabody technology (Fig. 8A; [31,33,34]). Globally, the structure of hSA3 is in agreement with the one of hSA1 solved by Khersonsky and colleagues and with the plasma-derived WT hSA structure, in the presence or absence of myristate, as supported by the root mean square deviation of 1.3 Å, calculated over all the Cα atoms, (Fig. 8C-D; PDB ID 2BXI, 2BXG, 7AAI; [15,24,35]).

**Fig. 8.**
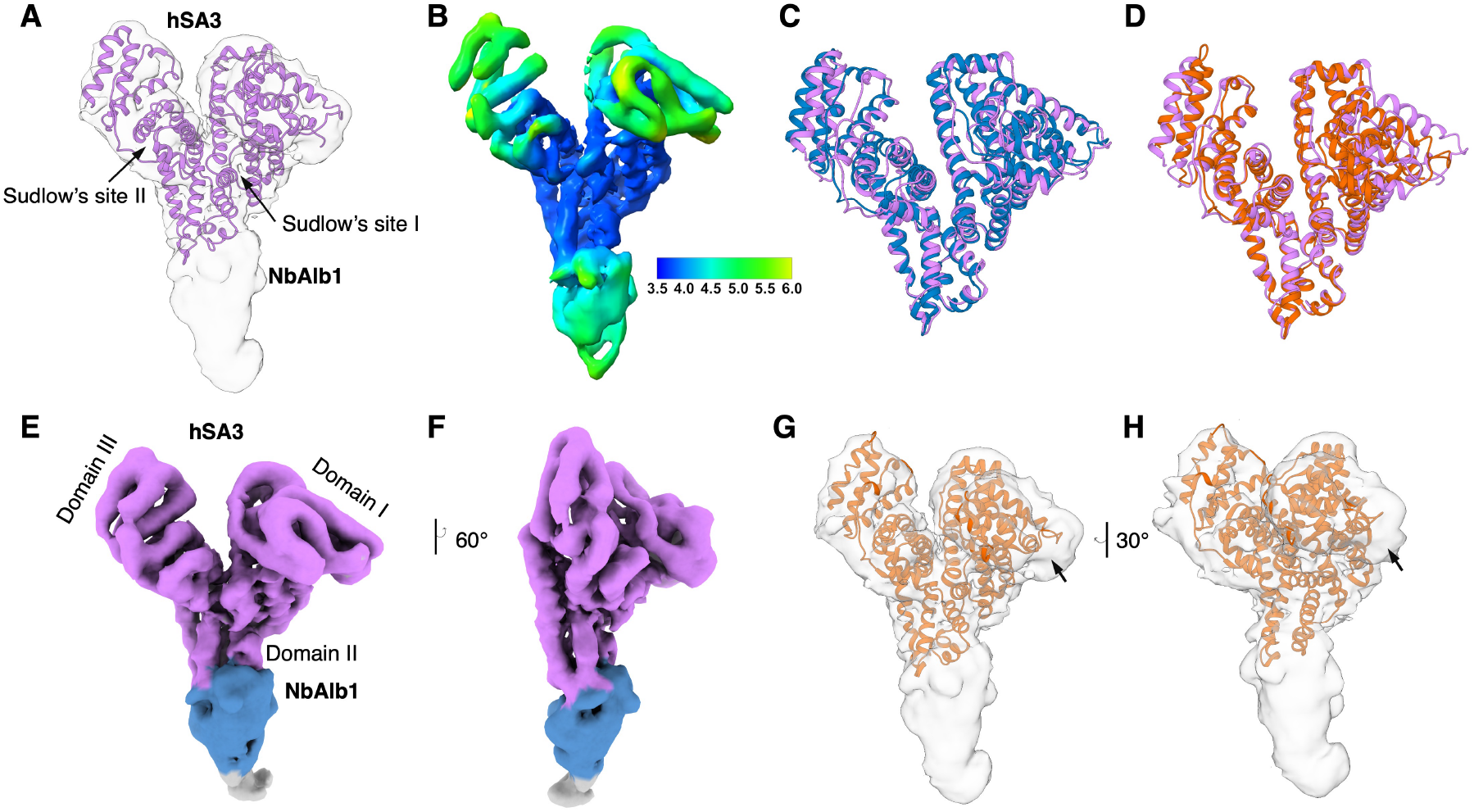
Cryo-EM structures of hSA3. (**A**) LocScale Cryo-EM density map of hSA3-MgbAlb1 with the fitted atomic model of hSA3; Sudlow site I and II are annotated according to the standard WT hSA nomenclature. (**B**) LocScale Cryo-EM density map of hSA3-MgbAlb1 colored by local resolution, highlighting that the core of hSA3 is resolved at higher resolution than the external domains. (**C**,**D**) hSA3 atomic model superimposed to WT hSA with and without myristic acid (PDB 2BXI and 2BXG) depicted in violet, blue, and orange respectively. (**E**) hSA3-MgbAlb1 LocScale Cryo-EM density map colored with the corresponding atomic model, (**F**) turned 60°. hSA3 density is depicted in violet, while NbAlb1 one in light-blue. hSA3 domains I, II, and III are annotated. (**G**,**H**) WT hSA compact conformation (without myristic acid; PDB code 2BXG) fitted into the hSA3-MgbAlb1 density. An arrow points to the unmodeled extra density, highlighting the open conformation of hSA3’s domain I.

Despite the global conservation, local differences are observed, especially in the conformation of domain I (Fig. 8G-H). Indeed, hSA3 domain I adopts the open conformation as the one observed for the plasma-derived WT hSA loaded with myristate (Fig. 8G-H; PDB ID 2BXI and 7AAI;[24,35]) and for the crystal structure of hSA1 (PDB ID 8A9Q; [15]). RMSD for domain I (residues 3–195) of hSA3 was 2.8 Å relative to myristate-bound WT hSA (PDB ID 2BXI), however increased to 6.0 Å when compared to the ligand-free WT structure (Fig. 8G-H). Although the resolution was not sufficiently detailed to distinguish the density of single fatty acid structures, such an open conformation could be the result of partial loading of fatty acids deriving from the culture medium and the bacterial expression host, analogously to that hypothesized in hSA1 crystallographic studies [15]. However, we cannot exclude that the mutations introduced in hSA3 *per se* favor this open flexible state, that may enhance pocket accessibility or alter the geometry of its binding sites, especially Sudlow site I, competent for warfarin binding capacity and largely affected by interdomain orientation and compactness ([24]; PDB ID 2BXI).

## 3. Conclusions

The growing demand for hSA across diverse fields, including pharmaceutical applications such as albumin-based vaccines and therapeutics, as well as in biochemical research and cell culture contexts, underlies the need for novel, recombinant, animal-free hSA variants.

In this study, we investigated the biophysical and structural properties of three highly mutated hSA variants; hSA1 (16 mutations), hSA2 (25 mutations) and hSA3 (73 mutations). These variants were designed *in silico* using the PROSS algorithm to make albumin expression in *Escherichia coli* available with high yield, thereby significantly reducing production costs.

Our experiments with warfarin and ibuprofen demonstrated that the introduced mutations tuned the binding affinities of the hSA variants to drugs. Although all designs exhibit the same order of magnitude in binding affinity for the ligands, the observed differences may be impactful for future developments of the carrier properties in the context of specific applications and, at the same time, open the possibility to explore novel delivery options.

SPR measurements demonstrated that the multiple sequence alignments performed by PROSS algorithm identified a set of mutations that enhanced protein stability and solubility without perturbing the interaction with hFcRn. Indeed, the affinity of the three engineered hSA variants to hFcRn are in line with that observed for WT hSA, exhibiting micromolar Kd at pH 5.5, reduced affinity at mildly acidic pH, and complete loss of binding at pH 7.4. The higher binding affinity for hFcRn observed for hSA1 and hSA2 at pH 6.0 may suggest that the variants can engage the receptor at an earlier stage in the endosomal compartments during cellular trafficking. This is interesting in light of the cellular data obtained using HERA, showing slightly more efficient recycling of hSA1 and hSA2 than that of hSA3. Overall, these findings demonstrate that the engineered variants can bind hFcRn in a pH-dependent manner and be recycled by the receptor. Studies in animal models are needed to evaluate whether the albumin engineering affects the pharmacokinetic properties of hSA in vivo. Cryo-EM analysis of the hypermutated hSA3, with the structure solved at an overall resolution of 3.9 Å, confirmed retention of the classical hSA global fold. The protein maintains its three subdomains arranged in the peculiar heart-shaped tertiary structure, in its open conformation. Furthermore, preliminary cytotoxicity assays confirmed the biocompatibility of the hSA variants at the tested concentration, suggesting their potential as an animal-free alternative in cell culture media. Future studies should focus on investigating their potential immunogenicity.

Through characterization and comprehensive structural assessments, we characterized a set of mutations that not only improve the stability and functionality of hSA but also provide insights into the mechanistic underpinnings of ligand interactions, recycling, and protein behavior. These findings pave the way for further advancements and applications of recombinant hSA variants in medicine and industry. Overall, we validate the structural integrity and functional fidelity of highly engineered hSAs, highlighting their potential as adaptable, animal-free platforms for advanced therapeutic development and wide industrial applications.

## 4. Material and Method

### 4.1 DNA constructs, Protein expression and purification

Commercial WT hSA was purchased from Albumin Bioscience (Albagen XL 9801). Commercial defatted WT hSA was purchased from Sigma-Aldrich (A1887). Plasmids pET29b with a C-terminal 6xHis tag containing the coding sequences of all the designed variants were kindly provided by Prof Sarel J. Fleishman [15]. hSA variants were purified from *Escherichia coli* SHuffle T7 strain as described by Khersonsky and colleagues, except for the introduction of a heat shock treatment before the immobilized metal affinity chromatography (IMAC) step. Supernatants from cell lysis were heated for 5’ at 10°C below each hSA variant melting temperature (*i*.*e*. 75°C, 80°C and 90°C, for hSA1, hSA2 and hSA3 respectively) and precipitated proteins were separated by centrifugation 4000xg, 20’, 20°C (Fig. S2-S3A; Eppendorf centrifuge 5810R, rotor F-34-6-38). Representative SDS-PAGE gel and chromatograms are represented in Fig. S2 [15]. Megabody Alb1 was purified as described by De Felice and colleagues [31]. pTT5 expression vector (NRCC) was used to produce a C-terminal FLAG/His tag construct of the β2-microglobulin domain fused to hFcRn α chain by a (G_4_S)_3_ linker (schFcRn; [15]), kindly provided by Prof. Dane Wittrup. FLAG/His-tagged schFcRn was expressed in FreeStyle Expi293FTM cells (Thermo Fisher Scientific) and purified with Ni-NTA resin (Cytiva) using 25 mM Tris, 150 mM NaCl supplemented with 500 mM imidazole. Eluted fractions were desalted in 25 mM Tris, 150 mM NaCl, pH 7.5, using a 5 mL HiTrap desalting column (Cytiva [36]). The purity and concentration of protein samples were evaluated by SDS-PAGE electrophoresis and UV absorption at 280 nm, respectively.

### 4.2 Dynamic light scattering

Dynamic Light Scattering (DLS) measurements were carried out using a Zetasizer Nano-S instrument (Malvern Instruments, UK) equipped with a He–Ne laser (4 mW, λ = 633 nm). Measurements were performed at a fixed backscattering angle of 173°, corresponding to the “non-invasive backscatter” (NIBS) configuration.

Samples were placed in disposable polystyrene cuvettes (pathlength 1 cm, volume 100 μL; Hellma, Switzerland) previously rinsed with filtered ultrapure water to minimize dust contamination. Prior to each measurement, the samples were filtered through a 0.22 μm PVDF syringe filter (Millipore) and equilibrated at 25 ± 0.1 °C for at least 10 minutes in the instrument cell holder to ensure thermal stability. For each sample, three consecutive measurements were recorded, each consisting of 10–15 subruns automatically optimized by the instrument’s software to achieve the best signal-to-noise ratio. The hydrodynamic diameter (Z-average, Dz) and polydispersity index (PDI) were calculated from the intensity autocorrelation function using the cumulant analysis method implemented in the Zetasizer Software (version 2.0.0.989).

All measurements were performed in triplicate, and the results are expressed as mean ± standard deviation (Fig. S3B-F). The viscosity and refractive index values of the solvent (e.g. PBS: η = 1.002 mPa·s, n = 1.330 at 25 °C) were entered in the software for data processing.

### 4.3 Isothermal Calorimetry analysis

Titrations of hSA variants with ligands were carried out at 310 K by using the MicroCal PEAQ-ITC instrument (Malvern Panalytical). hSA variants concentration was maintained to 70 µM or 120 µM in the cell and 1 mM Warfarin (Sigma-Aldrich A2250) or 4 mM Ibuprofen (Sigma-Aldrich I4883). Both hSA variants and ligands were dissolved in 50 mM NaPi, 100 mM NaCl, pH 7.4. Only in the case of Warfarin, 3% (v/v) DMSO was added to the buffer to improve its solubility. As a positive control, measurements were conducted with commercial WT hSA (Albagen XL 9801) and with commercial WT hSA, declared to be essentially defatted (Sigma-Aldrich, A1887). All measurements were performed in triplicate, and the results are expressed as mean ± standard deviation (Fig. S4-5).

In each experiment, an initial 0.4 µL injection (excluded from subsequent data analysis) was followed by 25 independent injections of 1.5 µL with a stirring rate of 750 rpm to guarantee rapid mixing [37]. A 120 s interval between injections was applied to ensure the equilibrium at each titration point. For each ligand a negative control was done measuring ligands against buffer and subtracted to the corresponding titrations to remove dilution heat contributions [37]. Data analysis was performed using the MicroCal PEAQ-ITC Evaluation Software (Malvern Panalytical, Malvern, UK). Integrated heat signals were fitted to “one set of sites” for all the measurements, values for the binding affinity constant (K_A_ = 1/K_D_) and the stoichiometry for each reaction were obtained from curve fitting [37].

### 4.4 Surface Plasmon Resonance

A Biacore T100 Instrument (GE Healthcare) was used with CM5 series S sensor chips (Cytiva) coupled with hFcRn^+k^ using amine coupling chemistry as described by the manufacturer, imposing a target level equal to 300 RU [27]. To assess the optimum ligand immobilization condition, a pH scouting was performed testing solutions of 10 mM ammonium acetate at pH equal to 3.5, 4.0, 4.5, 5.0, and 5.5. The coupling was performed by injecting 10 mg/mL of each protein into 10 mM sodium acetate, pH 4.5. Reference flow cells were blocked using 1 M ethanolamine (pH 8.5) to prevent the bulk effect of albumin. A running buffer of 25 mM MES, 150 mM NaCl, and 0.05% (v/v) Tween 20 (pH 5.5 or pH 7.4) was used as running buffer [38]. Single-cycle kinetics and affinity measurements were done by injecting serial dilutions of hSA1, hSA2, hSA3, WT hSA (Albagen XL 9801), defatted WT hSA (Sigma-Aldrich, A1887), from 0.3 µM to 30 µM with a flow rate of 50 mL/min, a contact time of 40 s and a dissociation step of 500 s done by using 25 mM MES, 150 mM NaCl, 0.05% (v/v) Tween 20, pH 5.5. Regeneration of the surfaces was obtained using injections of 30 mM Tris, 500 mM NaCl, 0.5 M MgCl_2_, pH 8.5 at 25°C, with a contact time of 30 s and flow rate of 30 mL/min.

Data were analyzed using the Biacore T100 Evaluation Software, zero adjusted and the reference cell value was subtracted and fitted by using the two-state reaction model, considering conformational change derived from the protonation of key histidine residues at acidic pH (Fig. 5). The measurements were done in triplicates (Fig. S6-8). The formula used by the software to evaluate the K_D_ value is the following: K_D_= kd1/ka1 * (kd2/(kd2+ka2)).

### 4.5 HERA and sample analysis

HERA was performed with HMEC-1 cells stably expressing hFcRn-eGFP (HMEC-1 hFcRn)), kindly gifted by Dr. Wayne I. Lencer (Boston Children’s Hospital, Harvard Medical School and Harvard Digestive Diseases Center, USA; [19,29,30]). The cells (1.5 × 10^5^ cells/mL) were seeded to 48-well plates in 250 μL growth medium per well containing MCDB-131 medium (Gibco) supplemented with 10% (v/v) heat-inactivated FBS (Gibco), 2 mM L-glutamine (Gibco), 1 µg/mL hydrocortisone (Sigma-Aldrich), 10 ng/mL recombinant mouse epidermal growth factor (Gibco), 25 U/mL penicillin and 25 µg/mL streptomycin (Gibco), 250 µg/mL geneticine selective antibiotic (G418 Sulfate; Gibco), and 5 µg/mL blasticidine S HCl (Gibco), and incubated for 22–24 h at 37 °C and 5% CO_2_. The growth medium was removed, and the cells were washed three times with 200 μL/well room-temperature HBSS (Gibco) and starved for 1 h in 250 μL/well HBSS. The buffer was gently removed and 800 nM of each protein of interest (hSA1, hSA2, hSA3,hSA isolated from human serum (Sigma-Aldrich, A1887), recombinant WT hSA and hSA QMP), diluted in 125 μL HBSS, was added to the respective wells in triplicate. Cells were incubated for 4 h to allow uptake of the proteins, followed by washing five times with 200 μL/well ice-cold HBSS. The wash buffer was removed, and the plate was placed on ice. The cells were lysed using 220 μL/well RIPA lysis buffer (Thermo Fisher) supplemented with 1X complete protease inhibitor cocktail (Roche). After 10 min of incubation with gentle tilting, the plate was centrifuged at 290-400 x g for 10 min at 4 °C. Cell lysates (“uptake” samples) were gently collected from the wells and stored at -20 °C until analysis. A separate, identically treated plate was washed as described after the 4 h-incubation, before 220 μL/well recycling medium (growth medium without FBS, G418 sulfate, or blasticidine, supplemented with 1X MEM non-essential amino acids solution; Gibco) was added. The cells were incubated for 18–20 h to allow cellular recycling, and the medium was then collected (“recycling” samples). All samples were stored at −20 °C until analysis.

A two-way anti-HSA ELISA was used to quantify the amount of protein in the samples from HERA, using a total volume of 100 µL/well in all steps, unless otherwise stated. Wells of clear flat-bottom 96-well EIA/RIA polystyrene microplates (Corning) were coated with 8⍰µg/mL polyclonal anti-hSA antibody from goat (Sigma-Aldrich, A1151) diluted in PBS overnight at 4 oC. The wells were then blocked with PBS supplemented with 2% (w/v) skimmed milk powder (PBSM) for 1 h at room temperature (RT) with gentle tilting. The buffer was removed from the wells, and samples from HERA diluted at DF1, DF2, or DF4 in PBSM supplemented with 0.005% (v/v) Tween 20 (Sigma-Aldrich; PBSTM) were added to the respective wells alongside a concentration gradient of each test article (0-8 nM, except 0-32 nM for hSA3). After 1 h of incubation at RT with gentle tilting, the wells were washed three times with 200 µL/well PBS supplemented with 0.005% (v/v) Tween 20 (Sigma-Aldrich). Protein was detected with 125 ng/mL ALP-conjugated polyclonal anti-hSA antibody from goat (Bethyl Laboratories Inc., A80-220AP) diluted in PBSTM and incubated 1 h at RT with gentle tilting. The wells were washed again as before, before 1 mg/mL p-nitrophenyl phosphatase substrate diluted in Pierce diethanolamine substrate buffer (Thermo Scientific) was added. A Sunrise microplate absorbance reader (TECAN) was used read absorbance in the wells at 405-620 nm.

Microsoft Excel and GraphPad Prism 10 were used for data analysis and presentation. To determine statistical differences, two-tailed unpaired Student’s *t*-tests were performed with a 95% confidence level and *p* < 0.05 defined as statistically significant.

### 4.6 Biocompatibility assay on human monocyte-derived macrophages

Buffy coats from healthy human donors were provided by the Blood Transfusion Centre of the Padova Hospital (Padova, Italy), complying with local regulations. Monocytes, isolated from buffy coats by Ficoll™ and Percoll™ gradient centrifugation, were resuspended in RPMI medium (ThermoFisher) at a concentration of 100 × 10^6^ cells/mL and further diluted to 8 × 106 cells/mL. Cells were seeded in 24-well plates (2 × 10^6^ cells/well) and incubated at 37 °C in a 5% CO_2_ for 1 h. Macrophages were then differentiated in fresh RPMI, supplemented with 10% (v/v) FBS and 100 ng/mL human macrophages colony stimulating factor (MCSF, Miltenyi, Germany). At day 7, the medium was changed to serum-free medium (X-VIVO^™^ 15, serum-free hematopoietic cell medium) and macrophages were incubated in triplicates with has formulations, LPS (ThermoFisher) and inorganic silica nanoparticles (Ludox®;[39,40]). Cytotoxic Ludox (1 mg/mL) was used as a positive control, LPS was used as a well-known activator of the immune system, inducing the clustering of macrophages, while serum free medium was used as negative control [39,41]. WT and hSA variants were tested at a final concentration of 0.4 mg/mL in line with previous data [15]. After 24 h, macrophages were detached by scraping, supplemented with PBS 1x supplemented with 1% (v/v) FBS to a final volume of 1 mL, centrifuged at 1200 rpm for 5 minutes at 4°C, and suspended in 300 mL of PBS 1x supplemented with 1% (v/v) FBS. After addition of PI (Sigma-Aldrich, 25 µg/mL), cell viability was measured by flow-cytometry using FACSDiva ® and analyzed using FACSDiva Software. A one-way ANOVA statistical test was performed to evaluate significant differences between the samples (GraphPad Prism 9).

### 4.7 hSA3 cryo-EM analysis

#### 4.7.1 Isolation of multi-protein complex hSA3-MgbAlb1 for cryo-EM studies

hSA3 and MgbAlb1c7HopQ were purified as described above [15,31], co-incubated at a final molar concentration of 50 µM and 100 µM, respectively, and then injected on a 200 10/300 GL SEC column (Cytiva), pre-equilibrated with 25 mM Tris, 150 mM NaCl pH 7.5 (Fig. S11A-B). Fractions corresponding to the hSA3-MgbAlb1c7HopQ binary complex were collected and deposited on grids for cryo-EM.

#### 4.7.2 Cryo-EM data collection

3.6 mL of freshly purified hSA3-MgbAlb1c7HopQ protein complex (0.3 mg/mL) were applied to a glow discharged Quantifoil R 1.2/1.3 Cu300 holey carbon grid. Excess sample was blotted away and plunge-frozen into liquid ethane using a Leica GP2 plunger (5 s blot time, at 25°C under 95% humidity) at the Kavli Institute of Nanoscience (TU Delft).

The grids were imaged on a 300 kV Titan Krios microscope (Thermo Fisher Scientific) at the Netherlands Centre for Electron Nanoscopy (NeCEN) facility with a K3 direct electron camera operated in the super-resolution mode at 0.418 Å per pixel. A total of 3732 movies were collected with 60 frames each, an exposure of 1 e/Å^−2^ per frame, and a defocus range from -2.4 µm to -1.6 µm (Fig. S11C; Table S3).

#### 4.7.3 Image Processing and 3D Reconstruction

Image processing pipeline was in CryoSPARC (v4.4; [42]). Motion correction was performed by Patch-Motion correction, a Fourier-crop factor of 0.5 was applied to bin the images to their physical pixel size of 0.836 Å and contrast transfer function (CTF) parameters were estimated using Patch-based CTF estimation [42]. 3032 micrographs were selected for the analysis and a random subset of 500 was selected for the initial processing steps. 327,743 particles were automatically picked by blob picker, extracted to a box size of 288 × 288 pixels, downsampled 4×, and subject to 2D alignment and classification. Particles from the best classes were selected for training the Topaz model [43], which was then used for automated particle picking from the full set of 3032 micrographs. 641,809 particles were extracted with 288 × 288 pixel box size and directly downsampled 4× into a 72×72 pixels. After several rounds of 2D classification, 316,549 particles were extracted to a box size of 288 × 288 pixels, 2× downsampled and used for an *ab-initio* reconstruction, then used as an input for a non-uniform refinement job that led to a consensus map of 4.2 Å overall resolution. Particles were further subjected to reference-based motion correction and then used as an input for a non-uniform refinement that led to a consensus map at 3.95 Å overall resolution [44,45]. Local amplitude scaling was performed using the model-free implementation of local sharpening with reference profiles in LocScale2 [46,47] with a cubic averaging window of 25 Å edge length and starting from the unfiltered half maps. The locally scaled map was used for display purposes; atomic model refinement and model-map Fourier shell correlation (FSC) were calculated using the original half maps (Fig. S12).

The complete pipeline used for the processing is summarized in Fig. S12, while representative 2D classes are depicted in Fig. S11D.

#### 4.7.4 Model building, refinement and structural analysis

A homology model for the atomic structure of hSA3 was generated by SwissModel using PDB ID 8A9Q (crystal structure of hSA1 in complex with warfarin; [15,48]) as a template. The atomic model of the binary complex, hSA3-NbAlb1 was obtained by combining the modelled hSA3 with NbAlb1, the latter extracted from the structure of wt hSA in complex with the same NbAl1 fusion construct, MgbAlb1c7HopQ (PDB ID 8OI2; [31]). The hSA3-NbAlb1 model was fitted in the cryo-EM density and refined by iterative cycles of automated real space refinement with Python-based Hierarchical Environment for Integrated Crystallography software (Phenix v1.20.1) with the simulated annealing and morphing option on [49–51]. A geometry minimization cycle was done to reduce the number of clashes (50 iterations, 1 macro cycle). Due to the low resolution of the map, the density corresponding to NbAlb1 was not interpreted.

Structural comparison analysis was performed using the General Efficient Structural Alignment of Macromolecular Targets (GESAMT) program of the CCP4 package [52]. All the RMSD calculations were computed by ChimeraX [53]. The hSA3 structural data have been validated and deposited in the Protein Data Bank archive with PDB code 9R8P and EM Data Resource with EMD code 53835.

### 4.8 Alphafold3 modelling of hSA2

Alphafold3 was used to model the structure of the hSA2 variant showed in Fig. 3 and 5 [54]. The protein sequence is reported in Fig. S6.

## Acknowledgments

We would like to thank the staff of the Netherlands Center for Electron Nanoscopy (NeCEN, Leiden, The Netherlands) for technical assistance during data collections. We thank Wiel Evers for technical assistance with the Geol microscope and Stefan T. Huber for technical assistance, data processing and model interpretation at the Kavli Institute of Nanoscience (Delft). We thank Olga Khersonsky and Sarel J. Fleishman for providing us the pET29b plasmids, technical support and critical reading of the manuscript. We acknowledge the financial support of the Italian Ministry of Education for supporting Sofia De Felice with a PhD scholarship. We thank the European Molecular Biology Organization (EMBO) for supporting Sofia De Felice with the EMBO Scientific Exchange Grant Initiative (grant number 10252). The work was partially supported by the Research Council of Norway through its Centres of Excellence scheme, project number 332727 (K.H.A., J.N., J.T.A.), and by grant 274993 (J.T.A., J.N.). The work was also supported by the South-Eastern Norway Regional Health Authority by grant 2024046 (K.H.A. and J.T.A.), and by grants 2018052 and 2019084 (J.T.A.).

## Author Contributions

**Sofia De Felice** writing, conceptualization, data curation, formal analysis, investigation, methodology, visualization, validation. **Christian Buratto** investigation, methodology, data curation **Alessia Savio** investigation, methodology, data curation, **Maria Morbidelli** conceptualization, methodology, data curation **Emanuele Papini** conceptualization, supervision, **Laura Acquasaliente** methodology, data curation, **Kristin Hovden Aaen** conceptualization, investigation, data curation, **Jeannette Nilsen** conceptualization, supervision, **Jan Terje Andersen** methodology, conceptualization, supervision, **Alessandro Angelini** methodology, conceptualization, resources, **Arjen J Jakobi** supervision, software, resources, **Laura Cendron** writing, conceptualization, data curation, project administration, resources, supervision, funding acquisition.

## Declaration of use of AI to improve readability and language

During the preparation of this work the author(s) used LucrezIA (developed by Unipd) in order to assist with language editing and refine grammar. After using this tool/service, the author(s) reviewed and edited the content as needed and take(s) full responsibility for the content of the publication.

