## supplementary figures for "Structural conservation and expanded functionality of hyper-stable human serum albumin variants"

```
hSA1      ------------------------DAHKSEVAHRFKDLGEENFKALVLIAFAQYLQQCPFEDLVKMVNEVTEFAKTCVADESAENCDKSLHTLFGDKLCTVATLRETYGEMADCCAKQEP 96
hSA2      ------------------------DAHKSEVAHRFKDLGEENFKALVLIAFAQYLQQCPFEDLVKMVNEVTEFAKTCVADESAENCDKSLHTLFGDKLCTVATLRETYGEMADCCAKQEP
hSA       MKWVTFISLLFLFSSAYSRGVFRRDAHKSEVAHRFKDLGEENFKALVLIAFAQYLQQCPFEDHVKLVNEVTEFAKTCVADESAENCDKSLHTLFGDKLCTVATLRETYGEMADCCAKQEP
hSA3      ------------------------DAHKSEVAHRFKDLGEENFKALVLIAFAQYLQKCPFEELVKMVKEVTEFAKKCVADETAENCDKSLHTLFGDKLCQVATLRETYGEMADCCAKQDP
mSA       MKWVTFLLLLFVSGSAFSRGVFRREAHKSEIAHRYNDLGEQHFKGLVLIAFSQYLQKCSYDEHAKLVQEVTDFAKTCVADESAANCDKSLHTLFGDKLCAIPNLRENYGELADCCTKQEP
                                  :*****:***::****::**.******:****:*.::: .*:*:***:***.*****:* ************** :..***.***:****:**:*

hSA1      EDLVKMVNEVTEFAKTCVADESAENCDKSLHTLFGDKLCTVATLRETYGEMADCCAKQEPERNECFLQHKDDNPNLPRLVRPEPDVMCTAFHDNEETFLKKYLYEIARRHPYFYAPELLY 156
hSA2      EDLVKMVNEVTEFAKTCVADESAENCDKSLHTLFGDKLCTVATLRETYGEMADCCAKQEPERNECFLQHKDDNPNLPRLVRPEPDVMCTAFHDNEETFLNKYLYEIARRHPYFYAPELLY
hSA       EDHVKLVNEVTEFAKTCVADESAENCDKSLHTLFGDKLCTVATLRETYGEMADCCAKQEPERNECFLQHKDDNPNLPRLVRPEVDVMCTAFHDNEETFLKKYLYEIARRHPYFYAPELLF
hSA3      EELVKMVKEVTEFAKKCVADETAENCDKSLHTLFGDKLCQVATLRETYGEMADCCAKQDPERHECFLQHKDDNPNLPRLERPEPDVMCKAFHDNEETFLNKYLYEIARRHPYFYAPELLY
mSA       DEHAKLVQEVTDFAKTCVADESAANCDKSLHTLFGDKLCAIPNLRENYGELADCCTKQEPERNECFLQHKDDNPSLPPFERPEAEAMCTSFKENPTTFMGHYLHEVARRHPYFYAPELLY
          :: .*:*:***:***.*****:* ************** :..***.***:****:**:***:**********.** : *** :.**.:*::*  **: :**:*:*************:

hSA1      FAKRYKAAFTECCQAADKAACLLPKLDELREEGKASSAKQRHKCAILQKFGERAFKAWAVARLSQRFPKAEFAEVSKLVTDLTKVHTECCHGDLLECADDRADLAKYICENQDSISSKLK 276
hSA2      FAKRYKAAFTECCQAADKAACLLPKLDELREEGKASSAKQRHKCAILQKFGERAFKAWAIARLSQRFPKAEFAEVSKLVTDLTKVHTECCHGDLLECMDDRADLAKYICENQDSISSKLK
hSA       FAKRYKAAFTECCQAADKAACLLPKLDELRDEGKASSAKQRLKCASLQKFGERAFKAWAVARLSQRFPKAEFAEVSKLVTDLTKVHTECCHGDLLECADDRADLAKYICENQDSISSKLK
hSA3      FAKRYKKAFTECCQAEDKAACLLPKLDALREEGKESSAKQRHKCAILQKFGERAFKAWAIARLSQRFPKAPFEEISKLVTDLTKVHKECCHGDMLECMDDRAKLAKYICENQDSISSKLK
mSA       YAEQYNEILTQCCAEADKESCLTPKLDGVKEKALVSSVRQRMKCSSMQKFGERAFKAWAVARLSQTFPNADFAEITKLATDLTKVNKECCHGDLLECADDRAELAKYMCENQATISSKLQ
          :*::*:  :*:**   ** :** **** ::::.  **.:** **: :************:***** **:* * *::**.******:.******:*** ****.****:**** :*****:

hSA1      ECCEKPLLEKSHCIAEVENDEMPADLPSLAADFIESKDVCKNYAEAKDVFLGMFLYEYARRHPDYSVVLLLRLAKTYETTLEKCCAAADPHECYSKVFDEFKPLIEEPQNLIKQNCELFE 396
hSA2      ECCEKPLLEKSHCIAEVENDEMPADLPSLAADFIESKDVCKNYAEAKDVFLGMFLYEYARRHPDYSVTLLLRLAKTYETTLEKCCAAADPHECYSKVFDEFKPLIEEPQNLIKQNCELFE
hSA       ECCEKPLLEKSHCIAEVENDEMPADLPSLAADFVESKDVCKNYAEAKDVFLGMFLYEYARRHPDYSVVLLLRLAKTYETTLEKCCAAADPHECYAKVFDEFKPLVEEPQNLIKQNCELFE
hSA3      ECCEKPLLERSHCIAEVENDDKPEDLPPLAADFVESKDVCKNYAEAKDVFLARFLYEYARRHPDYSVTLLLRIAKTYEDTLEKCCKAEDPHECYAKVEEEFKKLVEETQNLIKQNCELFE
mSA       TCCDKPLLKKAHCLSEVEHDTMPADLPAIAADFVEDQEVCKNYAEAKDVFLGTFLYEYSRRHPDYSVSLLLRLAKKYEATLEKCCAEANPPACYGTVLAEFQPLVEEPKNLVKTNCDLYE
           **:****:::**::***:*  * ***.:****:*.::*************. *****:******** ****:**.** ******   :*  **..*  **: *:**.:**:* **:*:*

hSA1      QLGEYKFQNALLIRYTKKVPQVSTPTLVEVARNLGKVGSKCCKHPEAKRMPCAEDYLSIVLNQLCVLHEKTPVSDRVTKCCTESLVNRRPCFSALEVDETYVPKEFNAETFTFHADICTL 516
hSA2      QLGEYKFQNALLIRYTKKVPQVSTPTLVEVARNLGKVGSKCCKLPEAKRMPCIEDYLSIVLNQLCVLHEKTPVNDRVTKCCTESLVNRRPCFSALEVDETYVPKEFNAETFTFHADICTL
hSA       QLGEYKFQNALLVRYTKKVPQVSTPTLVEVSRNLGKVGSKCCKHPEAKRMPCAEDYLSVVLNQLCVLHEKTPVSDRVTKCCTESLVNRRPCFSALEVDETYVPKEFNAETFTFHADICTL
hSA3      KLGEYYFQNALLIRYTKKMPQVPTDTLVELTRNLGKVGSKCCKLPEEKRLPCIEDYLSIVLNQLCVLHEKTPVNDRVTKCCTESLVNRRHCFSALEVDETYVPKEFNAETFTFHADLCTL
mSA       KLGEYGFQNAILVRYTQKAPQVSTPTLVEAARNLGRVGTKCCTLPEDQRLPCVEDYLSAILNRVCLLHEKTPVSEHVTKCCSGSLVERRPCFSALTVDETYVPKEFKAETFTFHSDICTL
          :**** ****:*:***:* ***.* **** :****:**:***. ** :*:** ***** :**::*:*******.::*****: ***:** ***** **********:*******:*:***

hSA1      SEEERQIKKQTALVELVKHKPKATKEQLKAVMDDFSAFVEKCCKADDKETCFAEEGKKLIAASQAALGL
hSA2      SEEERQIKKQTFLVELVKHKPKATKEQLKAIMDDFSAFVEKCCKADDKETCFAEEGKKLIAASQAALGL
hSA       SEKERQIKKQTALVELVKHKPKATKEQLKAVMDDFAAFVEKCCKADDKETCFAEEGKKLVAASQAALGL
hSA3      SEEERQIMKQKFLVELVKHKPKATEEQLKAVMDDFTAFVEKCCKAEDPETCFAEEGSKLIAKSQAALGL
mSA       PEKEKQIKKQTALAELVKHKPKATAEQLKTVMDDFAQFLDTCCKAADKDTCFSTEGPNLVTRCKDALA-
          .*:*:** **. *.********** ****::****: *::.**** * :***: ** :*:: .: **.
```

***Supplementary Figure 1. Multi sequences alignment of hSA variants, WT hSA and WT mSA performed by ClustalW.*** *Mutations introduced by PROSS design are highlighted in orange, pink and violet for hSA1, hSA2 and hSA3 respectively. The amino acid residues of WT mSA mutated with respect to WT hSA are highlighted in green.*

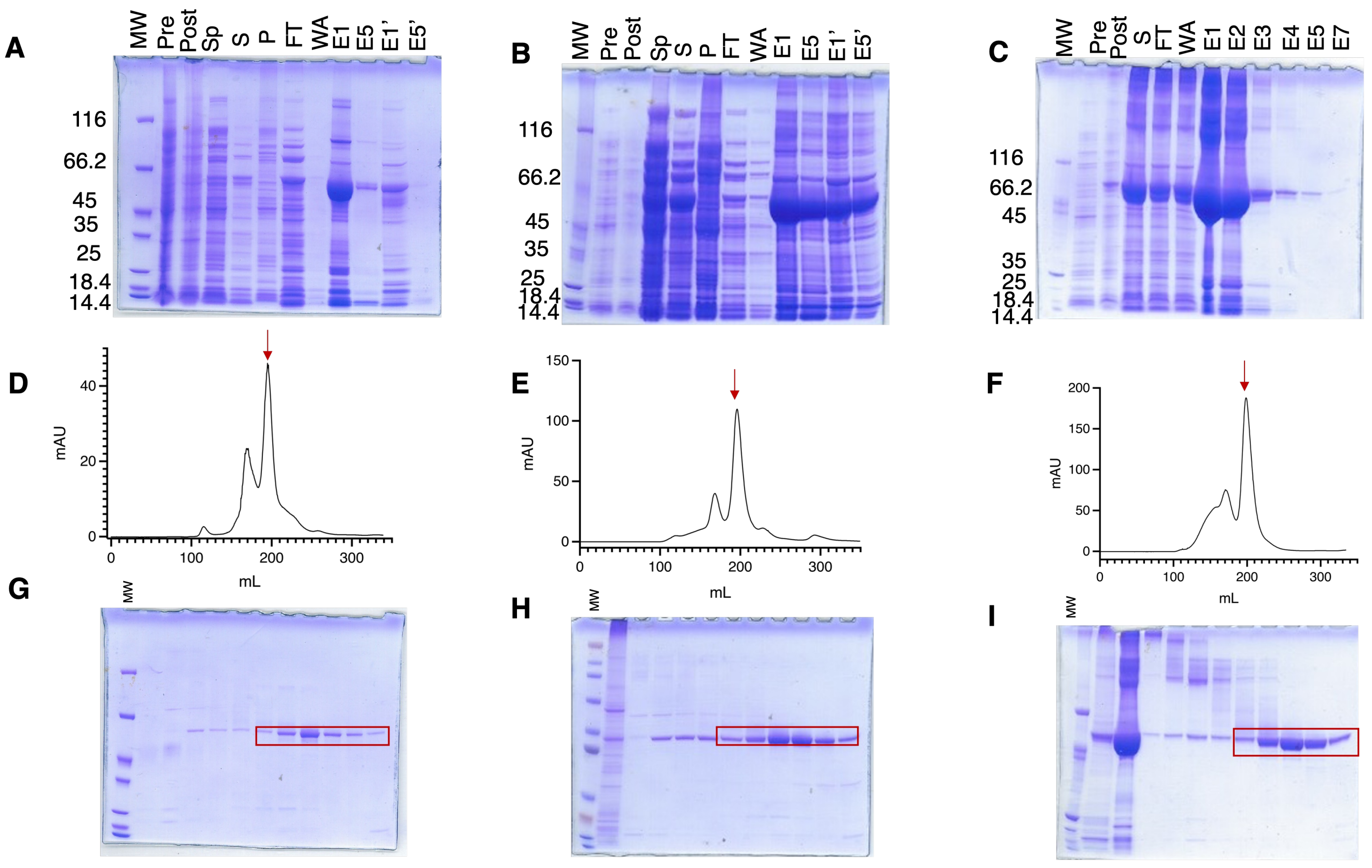


***Supplementary figure 2. Representative example of hSA variants purification.*** *(**A**,**B**,**C**) hSA variants purified by IMAC representative SDS-PAGE 4-12% run in MOPS, colored by Coomassie blue staining. FT: flow-through, WA: wash A, E n: Elution n. for hSA1,2 and 3. Pre (pre-induced culture); Post (post-induced culture); SP (supernatant before heat shock); S (supernatant after heat shock); P (pellet of precipitated protein after heat shock); FT (flow-through); WA (wash with PBS, pH 7.6); En (elution number with PBS+ 500 mM imidazole), En' (elution number with PBS+ 500 mM imidazole after a second FT loading). (**D**,**E**,**F**) SEC by using Superdex 200 126/60 column run in PBS pH 7.6 and (**G**,**H**,**I**) corresponding SDS-PAGE 4-12% run in MOPS and colored by Coomassie blue staining of hSA1,2,3. The peak corrisponding to hSA1,2,3 is pointed by a red arrow and boxed in red in the corresponding SDS-PAGE gel.*

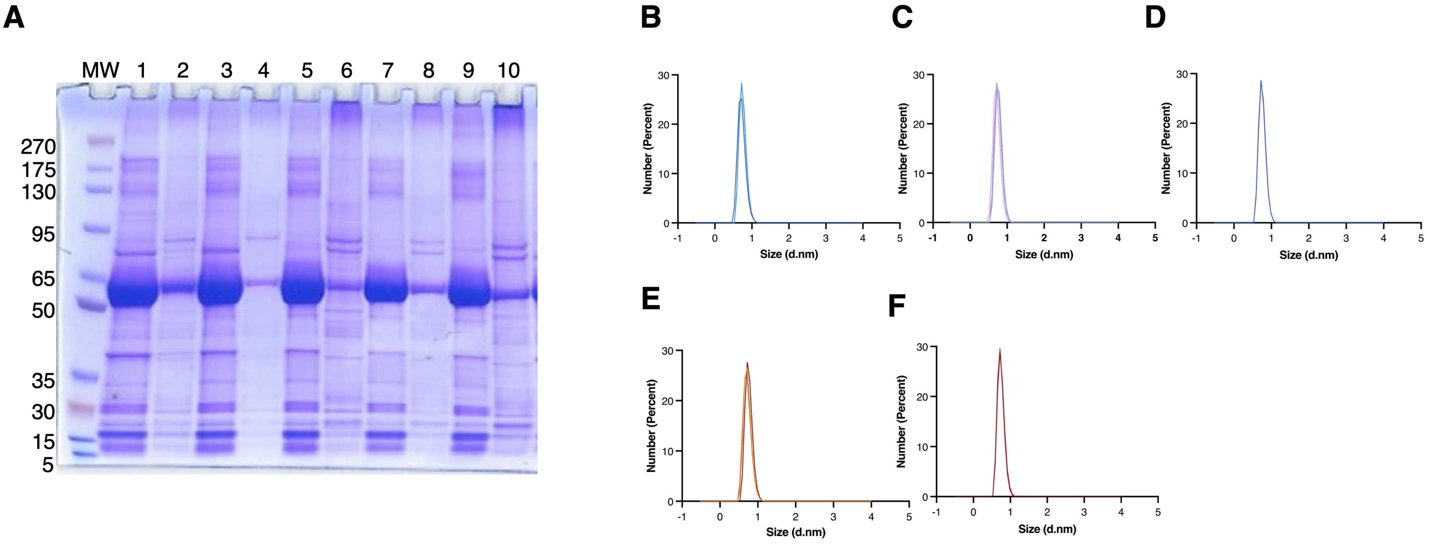


***Supplementary Figure 3. Heat shock treatment of hSA variants and DLS measurements. (A)*** *Representative SDS-PAGE gel 4-12%, run in MOPS, showing the heat-shock clarification preliminary test done for His-tagged hSA2. Supernatant after heating 3'/60°C (1), 5'/60°C (3), 3'/70°C (5), 5'/70°C (7), 5'/80°C (9) and corresponding pellet (2,4,6,8,10). Dynamic light scattering measurements triplicates for (**B**) defatted WT hSA, (**C**) WT hSA, (**D**) hSA1, (**E**) hSA2 and (**F**) hSA3.*

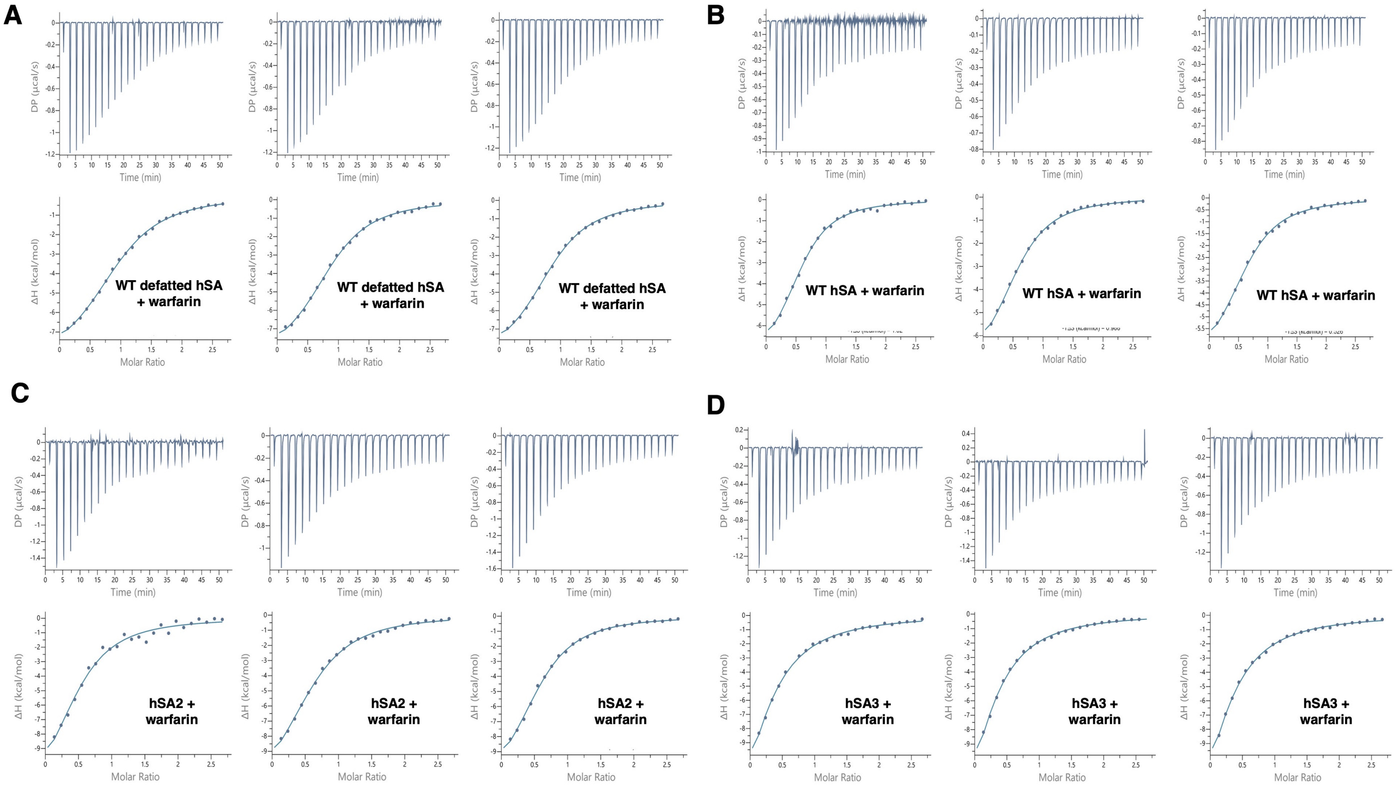


***Supplementary Figure 4. Triplicates of Warfarin binding to human serum albumin variants measured by isothermal titration calorimetry ITC.*** *(**A**) Warfarin binding to commercial defatted WT hSA, (**B**) WT commercial hSA, (**C**) hSA2 and (**D**) hSA3.*

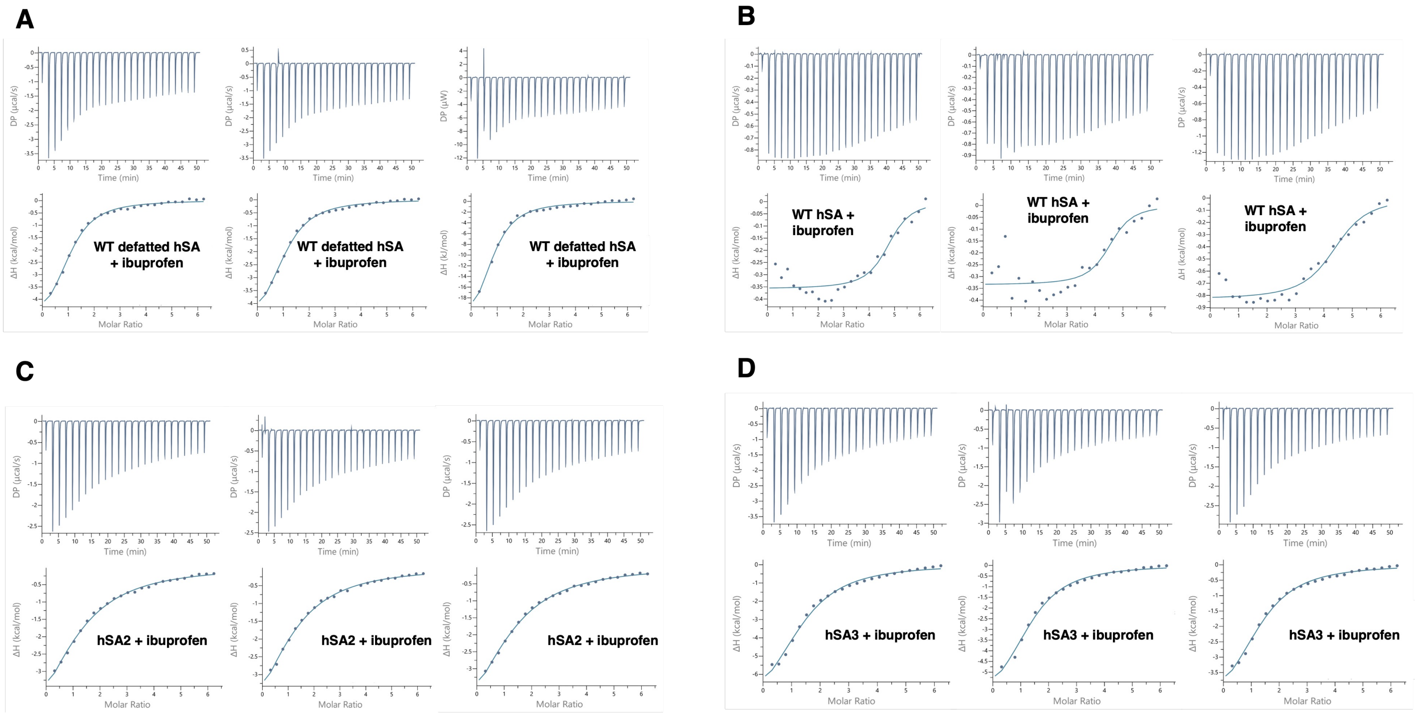


***Supplementary Figure 5. Triplicates of Ibuprofen binding to human serum albumin variants measured by isothermal titration calorimetry (ITC).*** *Ibuprofen binding to (**A**) commercial defatted WT hSA, (**B**) WT hSA, (**C**) hSA2 and (**D**) hSA3.*

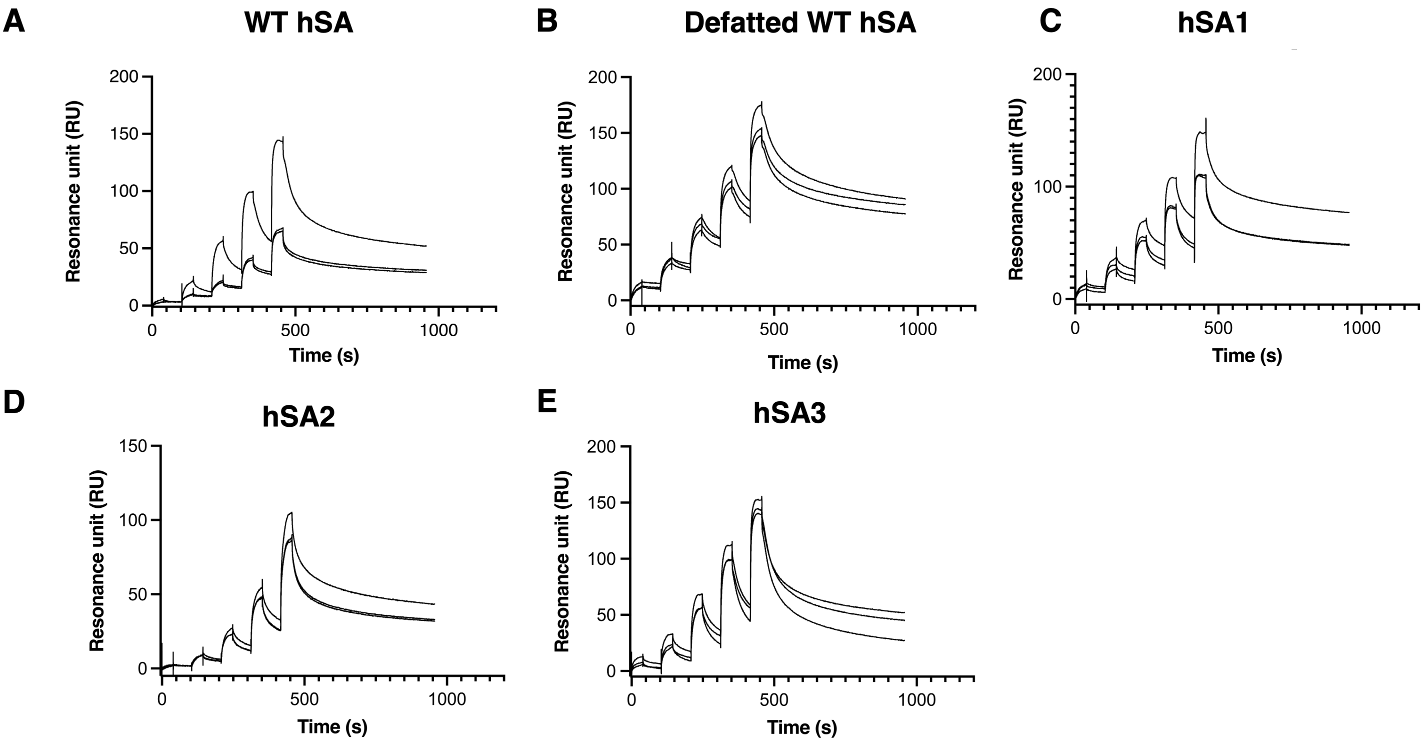


***Supplementary figure 6. Triplicates of single-cycle kinetics curves at pH 5.5****. Titrated concentration (0.37,1.1,3.3,10,30 µM) of (****A****) commercial WT hSA, (****B****) commercial defatted WT hSA, (****C****) hSA1, (****D****) hSA2, (****E****) hSA3 over the immobilized hFcRn (target level 300 RU).*

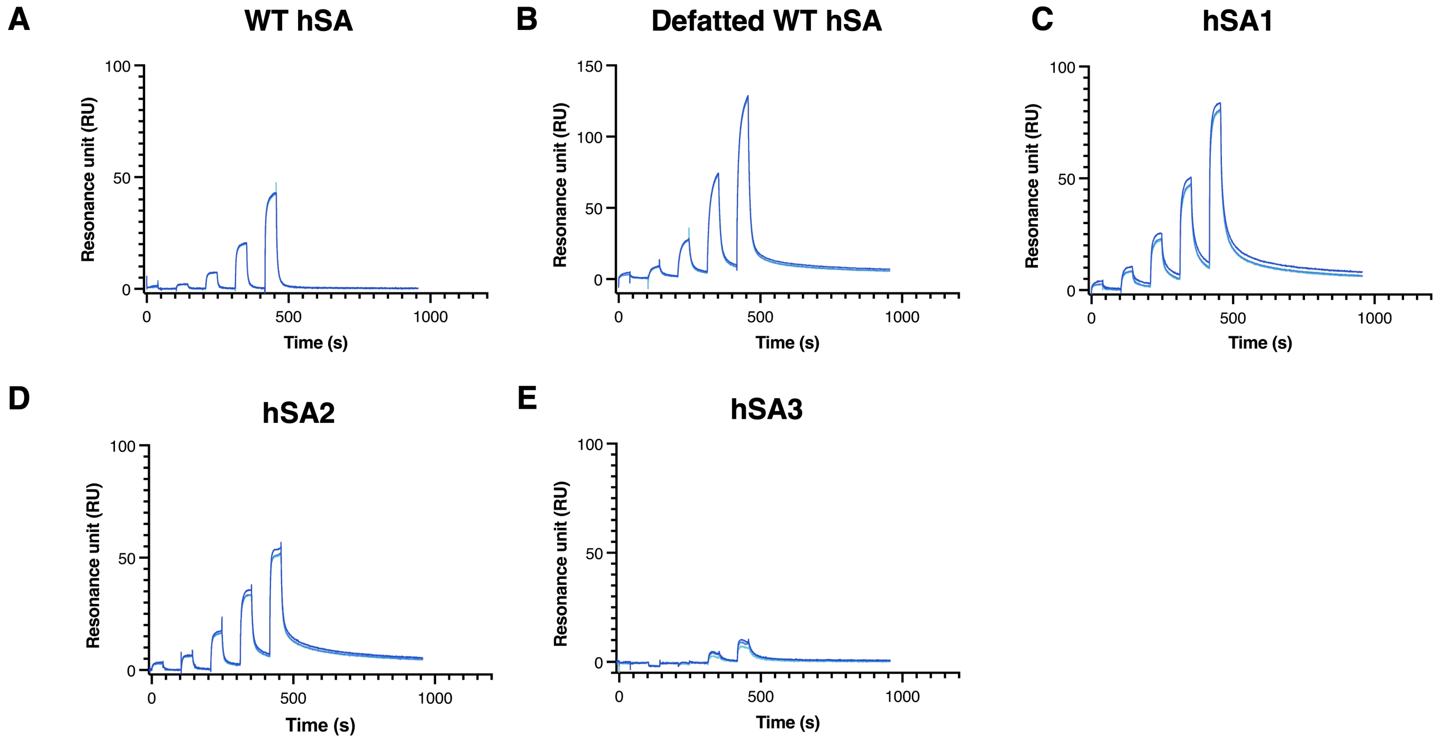


***Supplementary figure 7. Triplicates of single-cycle kinetics curves at pH 6***. *Titrated concentration (0.37,1.1,3.3,10,30* μM) *of (**A**) commercial WT hSA, (**B**) commercial defatted WT hSA, (**C**) hSA1, (**D**) hSA2, (**E**) hSA3 over the immobilized hFcRn (target level 300 RU).*

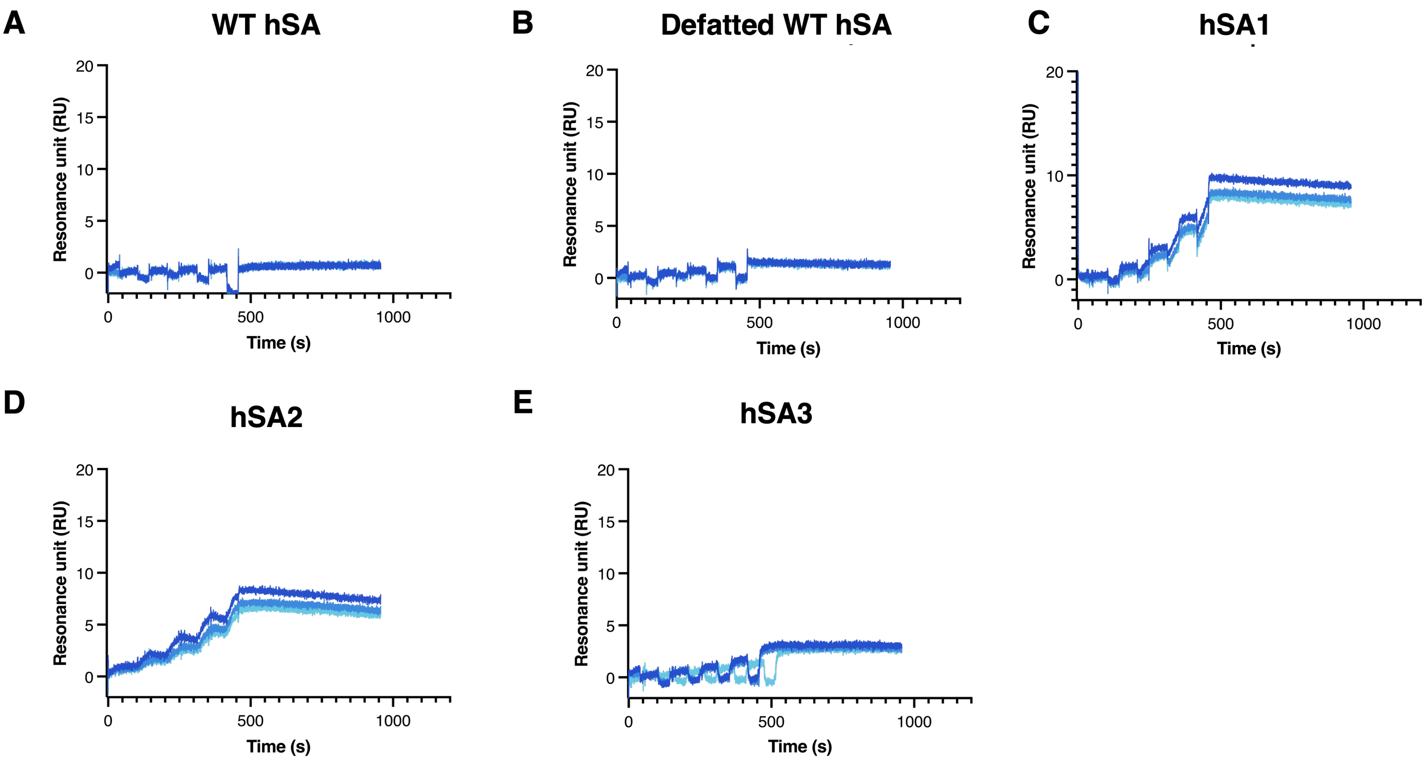


***Supplementary figure 8. Triplicates of single-cycle kinetics curves at pH 7.4**. Titrated concentration (0.37,1.1,3.3,10,30 µM) of (**A**) commercial WT hSA, (**B**) commercial defatted WT hSA, (**C**) hSA1, (**D**) hSA2, (**E**) hSA3 over the immobilized hFcRn (target level 300 RU).*

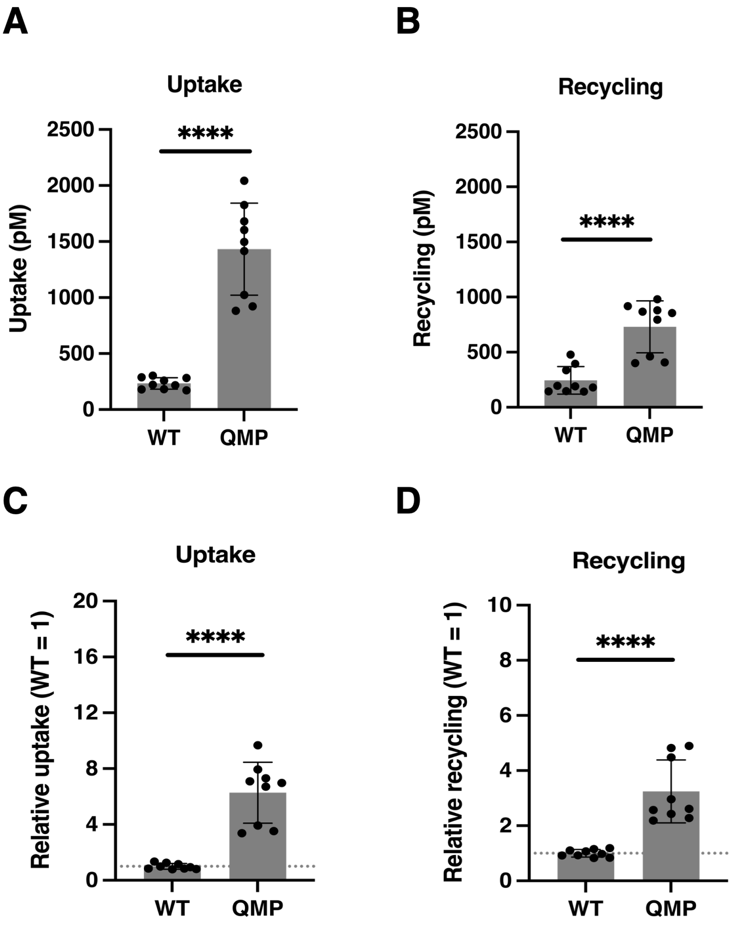


***Supplementary Figure 9. hFcRn-mediated cellular recycling of recombinant WT and QMP hSA.*** *Results from HERA with endothelial cells stably expressing hFcRn showing (**A, C**) cellular uptake of hSA variants and (**B, D**) hFcRn-mediated cellular recycling of hSA variants. Statistical significance was evaluated by Student's t tests (**** p < 0.001). Data are shown as protein concentration in sample or relative to that of recombinant WT hSA (WT = 1) and presented as the mean ± standard deviation (n = 9).*

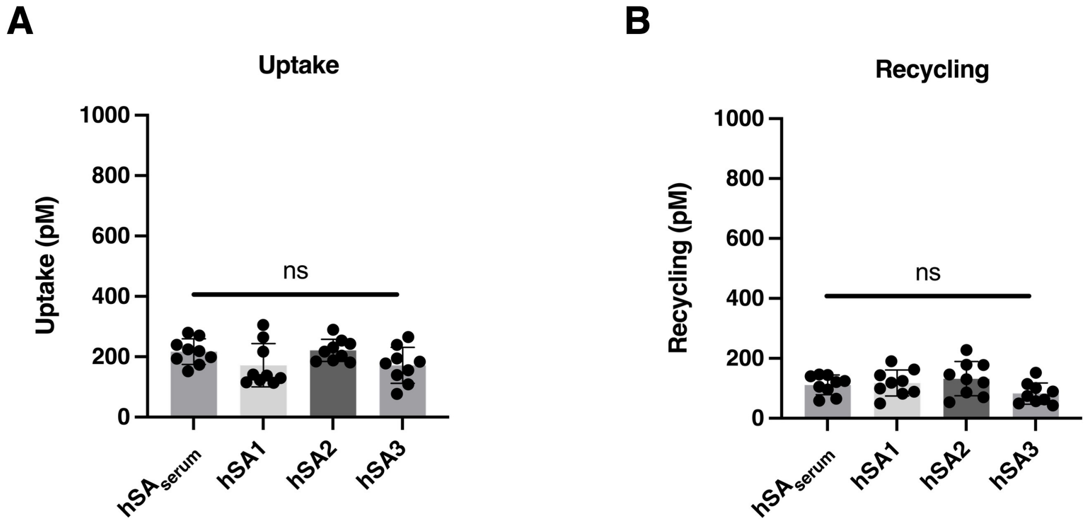


***Supplementary Figure 10. hFcRn-mediated cellular recycling and evaluation of biocompatibility of hSA variants***. *Results from HERA with endothelial cells stably expressing hFcRn showing (**A**) cellular uptake of hSA variants and (**B**) hFcRn-mediated cellular recycling of hSA variants. Statistical significance was evaluated by Student's t tests (ns, non-significant; $p > 0.05$). Data are presented as the mean* ± standard deviation *($n = 9$).*

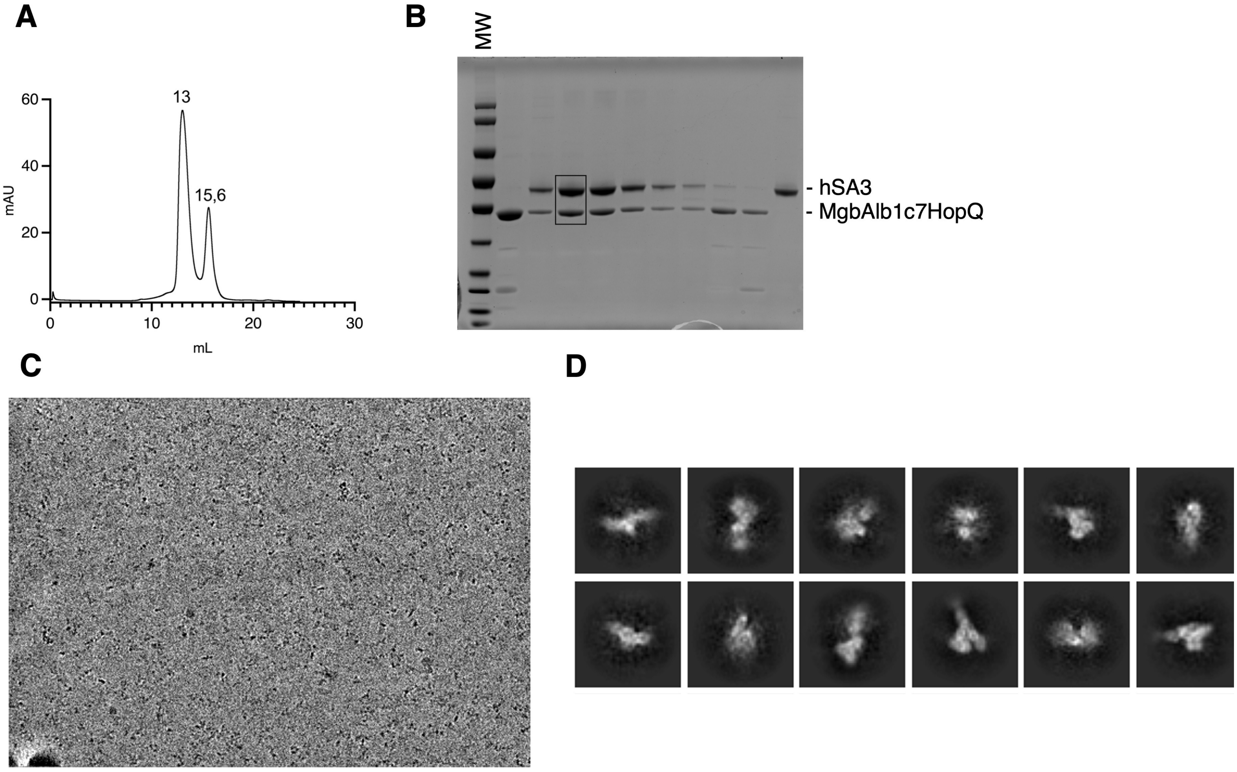


***Supplementary figure 11. Cryo-EM structure of hSA3-MgbAlb1: from sample preparation to data collection and analysis.*** *(**A**) hSA3-MgbAlb1 binary complex isolation, Superdex 200 10/300 analytical size exclusion chromatography of hSA3 in complex with MgbAlb1c7HopQ. The peak volume value is annotated above each curve. (**B**) SDS-PAGE gel 4-12% to check chromatograms fractions. (**C**) Cryo-EM representative micrograph of hSA3 in complex with MgbAlb1 and (**D**) 2D classes.*

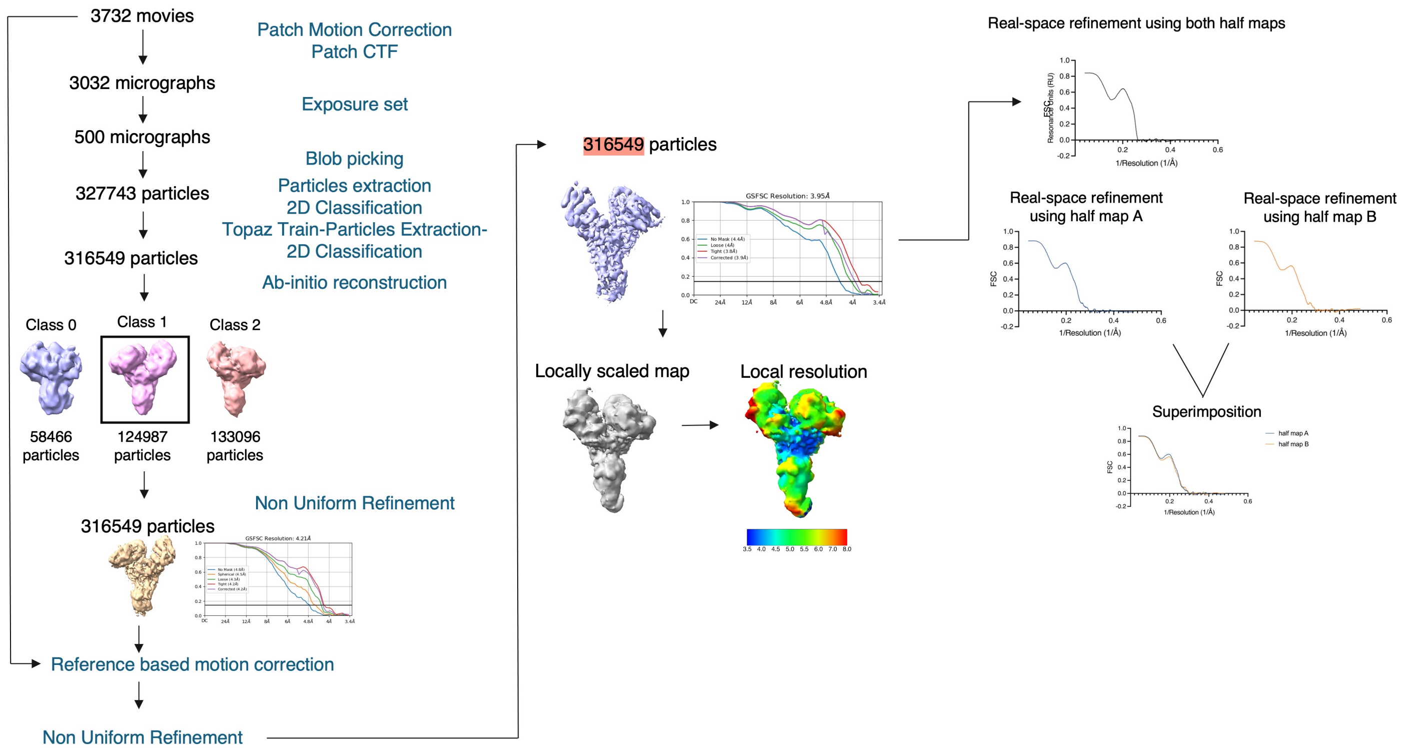


***Supplementary figure 12. hSA3-MgbAlb1 Cryo-EM analysis pipeline.*** *The data processing from movies alignment to the final non-uniform refinement and map post-processing is shown.*

***Supplementary Table 1*** *Residues nearby Sudlow site I (i.e., warfarin binding site) mutated in hSA variants. Residues in green are surface mutations, while the ones in yellow refer to mutations in ligand binding site. The ones in black are mutations neither in the ligand binding site nor on the surface.*

| Residue | WT hSA | hSA1 | hSA2 | hSA3 | Type of change |
|---|---|---|---|---|---|
| 156 | F | Y | Y | Y | apolar - polar (new H-bond) |
| 187 | D | E | E | E | negative - negative |
| 191 | A | A | A | E | non-polar - negative |
| 198 | L | H | H | H | non-polar- positive (new H-bond) |
| 202 | S | I | I | I | polar - non-polar |
| 216 | V | V | I | I | non-polar - non-polar |
| 243 | T | T | T | K | non-polar - positive |
| 254 | A | A | M | M | non-polar - non-polar |
| 286 | K | K | K | R | positive - positive |

***Supplementary Table 2.*** *Residues near the Sudlow site II (i.e., ibuprofen binding site) mutated in hSA variants. Residues in green are surface mutations, while the ones in yellow refer to mutations in ligand binding site. The ones in black are mutations neither in the ligand binding site nor on the surface.*

| Residue | WT hSA | hSA1 | hSA2 | hSA3 | Type of change |
|---|---|---|---|---|---|
| 344 | V | V | T | T | non-polar - polar |
| 349 | L | L | L | I | non-polar - non-polar |
| 384 | P | P | P | T | non-polar - polar |
| 397 | Q | Q | Q | K | polar - positive |
| 402 | K | K | K | Y | positive - polar |
| 409 | V | I | I | I | non-polar- non-polar |
| 415 | V | V | V | M | non-polar – non-polar |
| 419 | S | S | S | P | polar - non-polar |
| 421 | P | P | P | D | non-polar - negative |
| 426 | V | V | V | L | non-polar – non-polar |
| 427 | S | A | A | T | polar- non-polar |
| 440 | H | H | L | L | positive- non-polar |
| 446 | M | M | M | L | non-polar- non-polar |
| 449 | A | A | I | I | non-polar- non-polar |
| 455 | V | I | I | I | non-polar- non-polar |
| 470 | S | S | N | N | polar- polar |

***Supplementary Table 3.*** *Two state reaction model fits of single-cycle kinetics for triplicate measurements of hSA1,hSA2, hSA3, WT hSA and WT defatted hSA binding to immobilized hFcRn in 25 mM MES, 150 mM NaCl, 0.05% Tween-20, pH 5.5 and 6. Data were analyzed using T100 Evaluation software.*

| | Ka1 ($10^4$ 1/Ms) | Kd1 (1/s) | Ka2 (1/s) | Kd2 (1/s) | Kd2 (1/s) | KD (μM) | Rmax (RU) | Chi² ($RU^2$) |
|---|---|---|---|---|---|---|---|---|
| | | | | **pH 5.5** | | | | |
| **WT hSA (1)** | 1.31 | 0.1042 | 0.01686 | 0.001033 | 10.33 | 0.461 | 65.28 | 10.1 |
| **WT hSA (2)** | 0.3953 | 0.01932 | 0.01073 | 0.001079 | 10.79 | 0.447 | 55.99 | 4.12 |
| **WT hSA (3)** | 0.2604 | 0.0242 | 0.009408 | 0.001657 | 16.57 | 1.39 | 64.35 | 3.84 |
| **defatted WT hSA (1)** | 3.85 | 0.1548 | 0.01801 | 0.0008598 | 8.598 | 0.183 | 149.5 | 30.4 |
| **defatted WT hSA (2)** | 1.80 | 0.08176 | 0.01682 | 0.001019 | 10.19 | 0.260 | 142.8 | 26.7 |
| **defatted WT hSA (3)** | 0.6325 | 0.02135 | 0.01046 | 0.001091 | 10.91 | 0.319 | 168.8 | 39.9 |
| **hSA1(1)** | 3.30 | 0.07701 | 0.01138 | 9.23 10-4 | 9.23 | 0.175 | 93.42 | 15.7 |
| **hSA1(2)** | 2.31 | 0.08124 | 0.01104 | 9.53 10-4 | 9.53 | 0.279 | 101.7 | 15.2 |
| **hSA1(3)** | 2.92 | 0.1025 | 0.0137 | 10-4 | 8.30 | 0.201 | 142.3 | 27.2 |
| **hSA2 (1)** | 0.2480 | 0.02681 | 0.009687 | 0.001108 | 11.08 | 1.11 | 90.37 | 3.22 |
| **hSA2 (2)** | 0.2094 | 0.02082 | 0.008875 | 0.001055 | 10.55 | 1.06 | 90.88 | 7.2 |
| **hSA2 (3)** | 0.2018 | 0.02144 | 0.01052 | 0.001159 | 11.59 | 1.05 | 114 | 4.05 |
| **hSA3 (1)** | 0.6636 | 0.01994 | 0.00476 | 0.001258 | 12.58 | 0.628 | 139.2 | 19.4 |
| **hSA3 (2)** | 0.5245 | 0.02402 | 0.004881 | 0.002164 | 21.64 | 1.41 | 137.3 | 15 |
| **hSA3 (3)** | 0.4837 | 0.02105 | 0.006375 | 0.00114 | 11.4 | 0.660 | 139.1 | 14.4 |
| | | | | **pH 6** | | | | |
| **WT hSA (1)** | 0.7689 | 0.5319 | 0.02073 | 0.0714 | | 53.6 | 138 | 0.183 |
| **WT hSA (2)** | 0.8442 | 0.5773 | 0.02748 | 0.08427 | | 51.6 | 133.1 | 0.132 |
| **WT hSA (3)** | 0.7392 | 0.5104 | 0.02207 | 0.07367 | | 53.1 | 130.2 | 0.152 |
| **defatted WT hSA (1)** | 0.4037 | 0.101 | 0.002142 | 0.00211 | | 12.4 | 208.7 | 7.03 |
| **defatted WT hSA (2)** | 0.3874 | 0.104 | 0.002003 | 0.002322 | | 14.4 | 215.8 | 5.83 |
| **defatted WT hSA (3)** | 2.43 | 0.6778 | 0.00187 | 0.002263 | | 15.3 | 220.3 | 6.53 |
| **hSA1(1)** | 0.7847 | 0.05258 | 0.004116 | 0.002172 | | 2.32 | 65.5 | 1.77 |
| **hSA1(2)** | 0.6906 | 0.06037 | 0.004138 | 0.002522 | | 3.31 | 68.1 | 1.32 |
| **hSA1(3)** | 0.6603 | 0.06136 | 0.004139 | 0.002584 | | 3.57 | 69 | 1.32 |
| **hSA2 (1)** | 1.09 | 0.1387 | 0.0049 | 0.002491 | | 4.29 | 66.08 | 0.49 |
| **hSA2 (2)** | 1.05 | 0.1462 | 0.00464 | 0.002574 | | 4.97 | 66.11 | 0.424 |
| **hSA2 (3)** | 1.07 | 0.1464 | 0.00476 | 0.002646 | | 4.91 | 64.86 | 0.409 |
| **hSA3 (1)** | 0.3886 | 0.1224 | 0.004382 | 0.003468 | | 13.9 | 18.65 | 0.263 |
| **hSA3 (2)** | 0.3749 | 0.1289 | 0.005773 | 0.005073 | | 16.1 | 17.71 | 0.19 |
| **hSA3 (3)** | 0.2794 | 0.1556 | 0.006581 | 0.00782 | | 30.2 | 19.25 | 0.157 |

***Supplementary Table 4.*** *Cryo-EM data collection and model refinement statistics.*

| **Sample** | **hSA3** |
| --- | --- |
| EMDB ID | 53835 |
| PDB ID | 9R8P |
| **Data collection** | |
| Sample composition | hSA3-MgbAlb1c7HopQ |
| Microscope | Titan Krios |
| Magnification | 105000 |
| Voltage (kV) | 300 |
| Electron exposure ($e^-$/Å$^2$) | 60 |
| Exposure time (s) | 2.4 |
| Number of fractions | 60 |
| Number of movies | 3732 |
| Defocus range (µm) | -2.4/-1.6 |
| Pixel size (Å) | 0.418 |
| Detector | Gatan (K3) |
| Dose rate detector ($e^-$/Å$^2$/s) | 22.557 |
| **Data processing** | |
| Symmetry | / |
| Initial particles images (no.) | 327,743 |
| Final particles images (no) | 316,549 |
| Global map resolution (Å) | 4 |
| FSC threshold | 0.143 |
| Resolution range (local, Å) | 3.5-8 |
| **Model Refinement** | |
| CCmap_model | 0.66 |
| **Model composition** | |
| Non hydrogen atoms | 4715 |
| Protein residues | 4715 |
| **R.m.s deviations** | |
| Bond lengths (Å) | 0.003 |
| Angles (°) | 0.670 |
| **Ramachandran plot** | |
| Favored (%) | 94.31 |
| Allowed (%) | 5.69 |
| Outliers (%) | 0 |
| Clash score | 5 |
| Cβ outliers (%) | 0 |
