## supplementary tables for "Structural conservation and expanded functionality of hyper-stable human serum albumin variants"

**Supplementary Table 1** Residues nearby Sudlow site I (i.e., warfarin binding site) mutated in hSA variants. Residues in green are surface mutations, while the ones in yellow refer to mutations in ligand binding site. The ones in black are mutations neither in the ligand binding site nor on the surface.

| **Residue** | **WT hSA** | **hSA1** | **hSA2** | **hSA3** | **Type of change** |
| --- | --- | --- | --- | --- | --- |
| 156  187  191  198  202  216  243  254  286 | F  D  A  L  S  V  T  A  K | Y  E  A  H  I  V  T  A  K | Y  E  A  H  I  I  T  M  K | Y  E  E  H  I  I  K  M  R | apolar - polar (new H-bond)  negative - negative  non-polar - negative  non-polar- positive (new H-bond)  polar - non-polar  non-polar - non-polar  non-polar - positive  non-polar - non-polar  positive - positive |

**Supplementary Table 2.** Residues near the Sudlow site II (i.e., ibuprofen binding site) mutated in hSA variants. Residues in green are surface mutations, while the ones in yellow refer to mutations in ligand binding site. The ones in black are mutations neither in the ligand binding site nor on the surface.

| **Residue** | **WT hSA** | **hSA1** | **hSA2** | **hSA3** | **Type of change** |
| --- | --- | --- | --- | --- | --- |
| 344  349  384  397  402  409  415  419  421  426  427  440  446  449  455  470 | V  L  P  Q  K  V  V  S  P  V  S  H  M  A  V  S | V  L  P  Q  K  I  V  S  P  V  A  H  M  A  I  S | T  L  P  Q  K  I  V  S  P  V  A  L  M  I  I  N | T  I  T  K  Y  I  M  P  D  L  T  L  L  I  I  N | non-polar - polar  non-polar - non-polar  non-polar - polar  polar - positive  positive - polar  non-polar- non-polar  non-polar – non-polar  polar - non-polar  non-polar - negative  non-polar – non-polar  polar- non-polar  positive- non-polar  non-polar- non-polar  non-polar- non-polar  non-polar- non-polar  polar- polar |

**Supplementary Table 4.** Cryo-EM data collection and model refinement statistics.

| **Sample** | **hSA3** |
| --- | --- |
| EMDB ID  PDB ID  **Data collection**  Sample composition  Microscope  Magnification  Voltage (kV)  Electron exposure (e^–^/Å^2^)  Exposure time (s)  Number of fractions  Number of movies  Defocus range (μm)  Pixel size (Å)  Detector  Dose rate detector (e^–^/Å^2^/s)  **Data processing**  Symmetry  Initial particles images (no.)  Final particles images (no)  Global map resolution (Å)  FSC threshold  Resolution range (local, Å)  **Model Refinement**  CCmap_model  **Model composition**  Non hydrogen atoms  Protein residues  **R.m.s deviations**  Bond lengths (Å)  Angles (°)  **Ramachandran plot**  Favored (%)  Allowed (%)  Outliers (%)  Clash score  Cβ outliers (%) | 53835  9R8P  hSA3-MgbAlb1c7HopQ  Titan Krios  105000  300  60  2.4  60  3732  -2.4/-1.6  0.418  Gatan (K3)  22.557  /  327,743  316,549  4  0.143  3.5-8  0.66  4715  4715  0.003  0.670  94.31  5.69  0  5  0 |
